# *Cis* inhibition of co-expressed DIPs and Dprs shapes neural development

**DOI:** 10.1101/2024.03.04.583391

**Authors:** Nicholas C. Morano, Davys H. Lopez, Hagar Meltzer, Alina P. Sergeeva, Phinikoula S. Katsamba, Kevin D. Rostam, Himanshu Pawankumar Gupta, Jordan E. Becker, Bavat Bornstein, Filip Cosmanescu, Oren Schuldiner, Barry Honig, Richard S. Mann, Lawrence Shapiro

## Abstract

In *Drosophila*, two interacting adhesion protein families, Dprs and DIPs, coordinate the assembly of neural networks. While intercellular DIP/Dpr interactions have been well characterized, DIPs and Dprs are often co-expressed within the same cells, raising the question as to whether they also interact in *cis*. We show, in cultured cells and *in vivo,* that DIP-α and DIP-δ can interact in *cis* with their ligands, Dpr6/10 and Dpr12, respectively. When co-expressed in *cis* with their cognate partners, these Dprs regulate the extent of *trans* binding, presumably through competitive *cis* interactions. We demonstrate the neurodevelopmental effects of *cis* inhibition in fly motor neurons and in the mushroom body. We further show that a long disordered region of DIP-α at the C-terminus is required for *cis* but not *trans* interactions, likely because it alleviates geometric constraints on *cis* binding. Thus, the balance between *cis* and *trans* interactions plays a role in controlling neural development.

## INTRODUCTION

The assembly of complex neural networks requires precise wiring of many types of neurons to ensure the formation of functional circuits. In the developing nervous systems of both vertebrates and invertebrates, neurons express a wide diversity of cell adhesion molecules (CAMs), which coordinate neural network development by mediating cell-cell recognition^1,2^. In *Drosophila*, two interacting adhesion protein families, the 21-member Dpr (Defective proboscis response) proteins and the 11-member DIPs (Dpr interacting proteins)^3^ are required for the development of several neural networks^4–7^. Both DIPs and Dprs are immunoglobulin superfamily (IgSF) proteins, containing 3 and 2 Ig domains, respectively. DIP::Dpr interactions were initially identified in a high-throughput screen^8^, and have since been extensively characterized using structural and biophysical approaches ^5–11^. Moreover, multiple studies have demonstrated that specific subsets of DIPs and Dprs are expressed in different neurons throughout the developing *Drosophila* nervous system. *Trans* (cell-to-cell) interactions between specific cognate pairs of DIP and Dpr family members specify neuronal and neuromuscular connectivity in the developing *Drosophila* nervous system^3,4,7,9,12^. For example, in the medulla of the fly’s visual system, DIPs are expressed in a layer-specific manner and interaction with their cognate Dprs is required for incoming axons to form layer-specific synapses^13^. More recently, we identified a specific *trans*-interaction between two types of neurons in the developing fly mushroom body as essential for target specificity^14^.

While the DIPs tend to have specific expression patterns, Dprs are often widely expressed throughout the nervous system, resulting in the co-expression of interacting family members in the same cells^12,14–16^. Although the importance of *trans* DIP::Dpr interactions and homophilic *cis* DIP interactions is well-characterized^17^, the functional role of cognate partners that are co-expressed within the same cell remains elusive.

While *cis* interactions are prevalent among CAMs, they often employ separate interfaces to facilitate *cis* and *trans* interactions^18–20^. For example, for the clustered protocadherins, which regulate neuronal self-avoidance, heterophilic *cis* interactions and homophilic *trans* interactions are mediated through separate interfaces. The classical cadherins such as N-Cadherin and E-Cadherin also have distinct *cis* and *trans* interfaces with weak *cis* interactions enabling the formation of two-dimensional molecular lattices^21,22^. Conversely, the Sidekicks (Sdks), which like the DIPs and Dprs are members of the IgSF, interact both in *cis* and in *trans* through the same interface^23^. We show below that this is the case for DIPs and Dprs as well. Although *cis* interactions between CAMs are widespread, the mechanisms of *cis* binding and biological functions of *cis* interactions are less well understood.

Here, we test the idea that *cis* interactions between cognate DIP::Dpr pairs play a role in neural connectivity by focusing on two sets of interacting DIP::Dpr proteins: DIP-α with Dprs 6 and 10, and DIP-δ with Dpr12. We use flow cytometry-based assays to demonstrate that co-expression of DIP-α with Dprs 6 and 10 effectively blocks binding in *trans*, arguing that binding in *cis* competes with *trans* binding. We demonstrate similar findings for DIP-δ and Dpr12, suggesting a potential role for *cis* inhibition for this cognate pair as well. In contrast, we find that DIP-β does not exhibit *cis* inhibition. Based on sequence analysis of DIPs α, β and δ, we propose a model for DIP::Dpr *cis* inhibition in which the length of the linker between the C-terminal Ig domain and the glycosylphosphatidylinositol (GPI) cell membrane anchor of the DIP is a crucial determinant of *cis* binding. We then test, both in cells and in vivo, whether the long flexible linkers are needed to overcome the geometric constraints that would otherwise prevent binding between cognate DIPs and Dprs when present on the same membrane.

We use two *in vivo* systems to assess the role of *cis* inhibition during *Drosophila* development: the mushroom body (MB) and the neuromuscular junction (NMJ). In the MB of the developing pupal brain, γ-Kenyon cells (γ-KCs) express Dpr12, which interacts in *trans* with its sole binding partner, DIP-δ. We demonstrate that mis-expressing DIP-δ in γ-KCs results in circuit malformations that phenocopy Dpr12 loss-of-function, consistent with the idea that DIP-δ binds Dpr12 in *cis* within γ-KCs, reducing its availability to interact with DIP-δ in *trans*. In the developing legs of the fly, a *trans* interaction between DIP-α, expressed in only three motor neurons (MNs), and Dpr10, expressed broadly in leg muscles, is required for the terminal branching and maintenance of MN–muscle contacts^12^. We use ectopic expression, knockdown experiments, and CRISPR/Cas9 genetic manipulations to demonstrate that co-expression of Dpr6/10 with DIP-α in MNs modulates the ability of these molecules to interact in *trans*, and can lead to significant alterations of neuronal morphology. Taken together, both sets of *in vivo* experiments indicate that the correct balance between *cis* and *trans* interactions is critical for the correct assembly of neural circuits.

## RESULTS

### Quantification of cell surface DIPs and Dprs using flow cytometry

We developed a flow cytometry-based platform to detect cell surface protein interactions of DIPs and Dprs, which had previously been used to investigate members of the human IgSF ^24,25^. Typically, an interacting receptor::ligand pair is tagged with different-colored fluorescent proteins (GFP and mCherry), expressed in HEK293 cells, and incubated together. The interaction of protein pairs can then be detected using flow cytometry by analyzing the number of events positive for both GFP and mCherry (**Figure 1A**). Because the cell surface attachment of DIPs and Dprs requires GPI anchors, we avoided adding tags that could disrupt membrane attachment. Instead, all DIPs and Dprs were subcloned into a mammalian expression vector containing an Internal Ribosome Entry Site (IRES) with either mCherry (DIPs) or GFP (Dprs) at the second ribosomal entry site. DIP/Dpr constructs were individually transfected into HEK293 cells and all showed expression of GFP and mCherry (**Supplementary Figure S1**). Cells transfected with DIPs were then incubated in 96 well plates with cells transfected with Dprs, and the aggregation of DIP::Dpr pairs was assessed using flow cytometry (**Figure 1B-F**).

**Figure 1:**
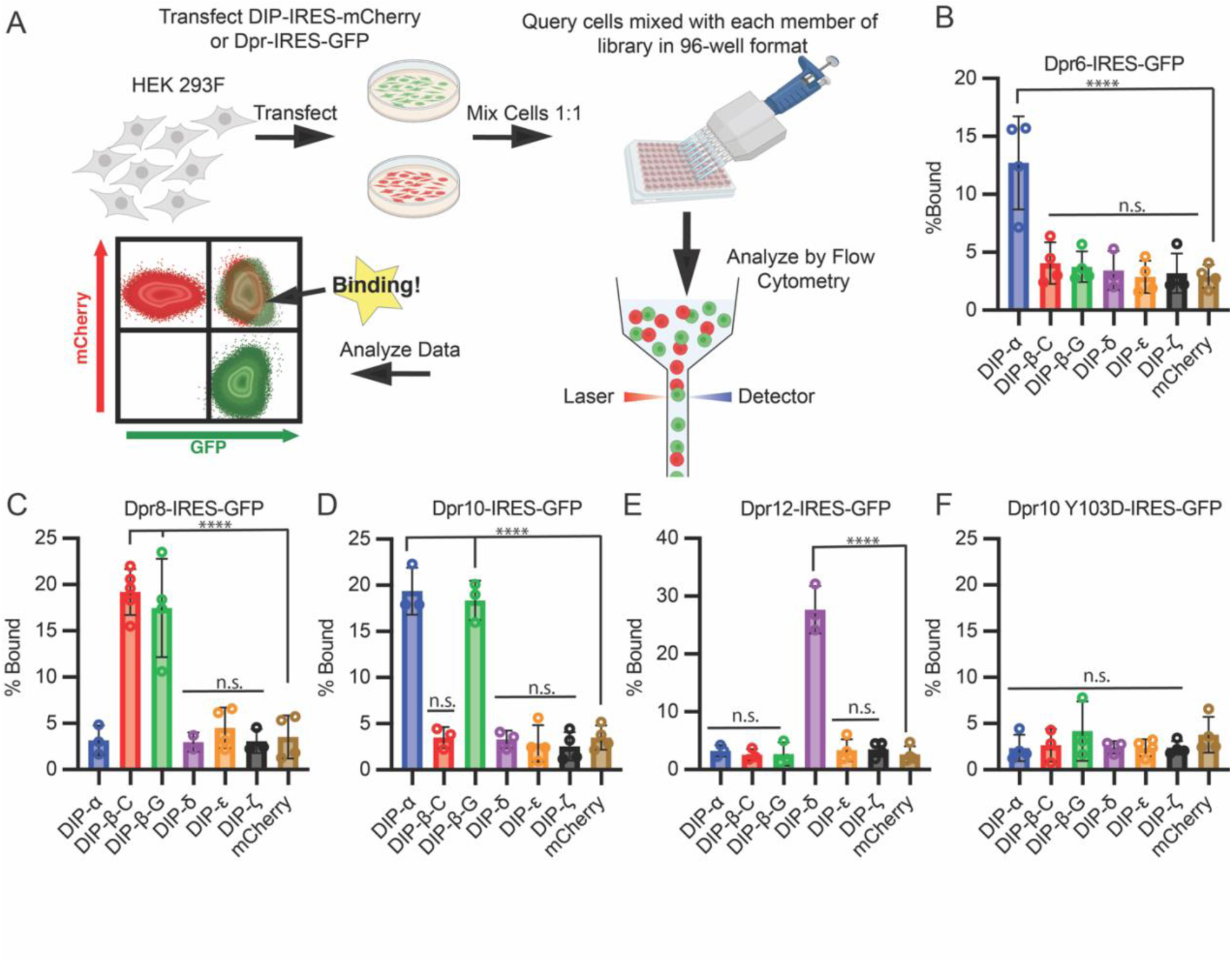
Flow cytometry based cell adhesion assay detects high affinity DIP::Dpr Interactions. **(A)** Overview schematic of flow-based cell adhesion platform. **(B-F)** Proof of concept experiment demonstrating that full-length DIPs and Dprs can be expressed in HEK293 Freestyle cells, and that adhesion can be measured by flow cytometry. An initial set of Dpr6 **(B)**, Dpr8 **(C)**, Dpr10 **(D)**, and Dpr12 **(E)** were expressed with an IRES GFP tag and queried against a set of DIP-IRES mCherry constructs. Adhesion was detected only between DIP/Dpr pairs with high affinity molecular interactions. A negative control, Dpr10 with Y103D mutation introduced, did not interact with DIP-α **(F)** as expected.

We generated a collection of expression vectors encompassing a broad range of interaction affinities composed of DIP-α, β, δ, ε, and ζ, and Dprs 6, 8, 10, and 12. We also included the DIP-β-G isoform, which has an 11 amino acid deletion in the first Ig domain relative to DIP-β-C as well as two isoforms of Dpr10, denoted here as Dpr10-A and Dpr10-D, with isoform A binding somewhat stronger to DIP-β-G than isoform C (**Supplementary Figure S2**). Unless otherwise specified DIP-β will refer to DIP-β-C and Dpr10 will refer to Dpr10-A. We also report experiments on the Y103D mutation of Dpr10 which ablates DIP::Dpr interactions^9^.

**Supplementary Data File 1** reports equilibrium binding constants (K_D_s) determined using surface plasmon resonance (SPR for DIP::Dpr pairs used in this study. Most of these data have been published previously ^3,9^, although measurements on DIP-β-G are new (**Supplementary Figure S2)**. All interactions with lower K_D_s^4,8,9^ listed in **Supplementary Data File 1** were detected by flow cytometry, while interactions with higher K_D_s were not (**Figure 1B-F**). In particular, Dpr6 bound to DIP-α but not other DIPs (**Figure 1B**), while Dpr8 specifically bound to DIP-β isoforms C and G (**Figure 1C**), Dpr-10 bound to DIP-α and also DIP-β-G (but not DIP-β-C, **Figure 1D**) and finally, Dpr12 specifically bound to DIP-δ (**Figure 1E**). Dpr10 Y103D, which ablates its ability to interact with DIP-α and DIP-β, also failed to show binding using this assay (**Figure 1F**). Taken together, these results confirm the cell-based expression of these DIPs and Dprs, and validate a flow cytometry platform for quantitatively examining the binding properties of DIPs and Dprs.

### Co-expression of DIP-α and DIP-**δ**, but not DIP-β, with their cognate Dprs inhibits interactions with high affinity ligands

To perform cell-protein binding experiments, Dpr10-Fc was purified as an Fc fusion protein, which dimerizes due to the Fc domain. This artificial dimer (**Figure 2A**) binds robustly to cells expressing DIP-α (**Supplementary Figure S4A**). In order to test whether co-expressed DIPs and Dprs can interfere with this binding, Dpr10-Fc was titrated from 1-600 nM onto cells expressing DIP-α, co-expressing DIP-α and Dpr10, or control cells (transfected with mCherry). We obtained robust binding curves for both A and D isoforms of Dpr10 (see **Supplementary Figure S2-3** for related SPR data) with DIP-α expressing cells. No binding was detected to cells co-expressing DIP-α and Dpr10 or control cells, demonstrating that expression of DIP-α and Dpr10 in *cis* inhibited binding of DIP-α to recombinant Dpr10-Fc (**Figure 2B, Supplementary Figure S4A-B**). Dpr10-Fc was also titrated against cells co-expressing DIP-α and Dpr6 (**Figure 2C**). Binding was detected to cells expressing DIP-α, but not to cells co-expressing DIP-α and Dpr6. These data suggest that co-expression (in *cis*) of DIP-α with either Dpr6 or Dpr10 inhibits trans-binding of DIP-α with Dpr10. Likewise, Dpr12-Fc binds to cells expressing DIP-δ, but not to cells co-expressing DIP-δ and Dpr12, which also indicates *cis* inhibition (**Figure 2D, Supplementary Figure S4C-D**).

**Figure 2:**
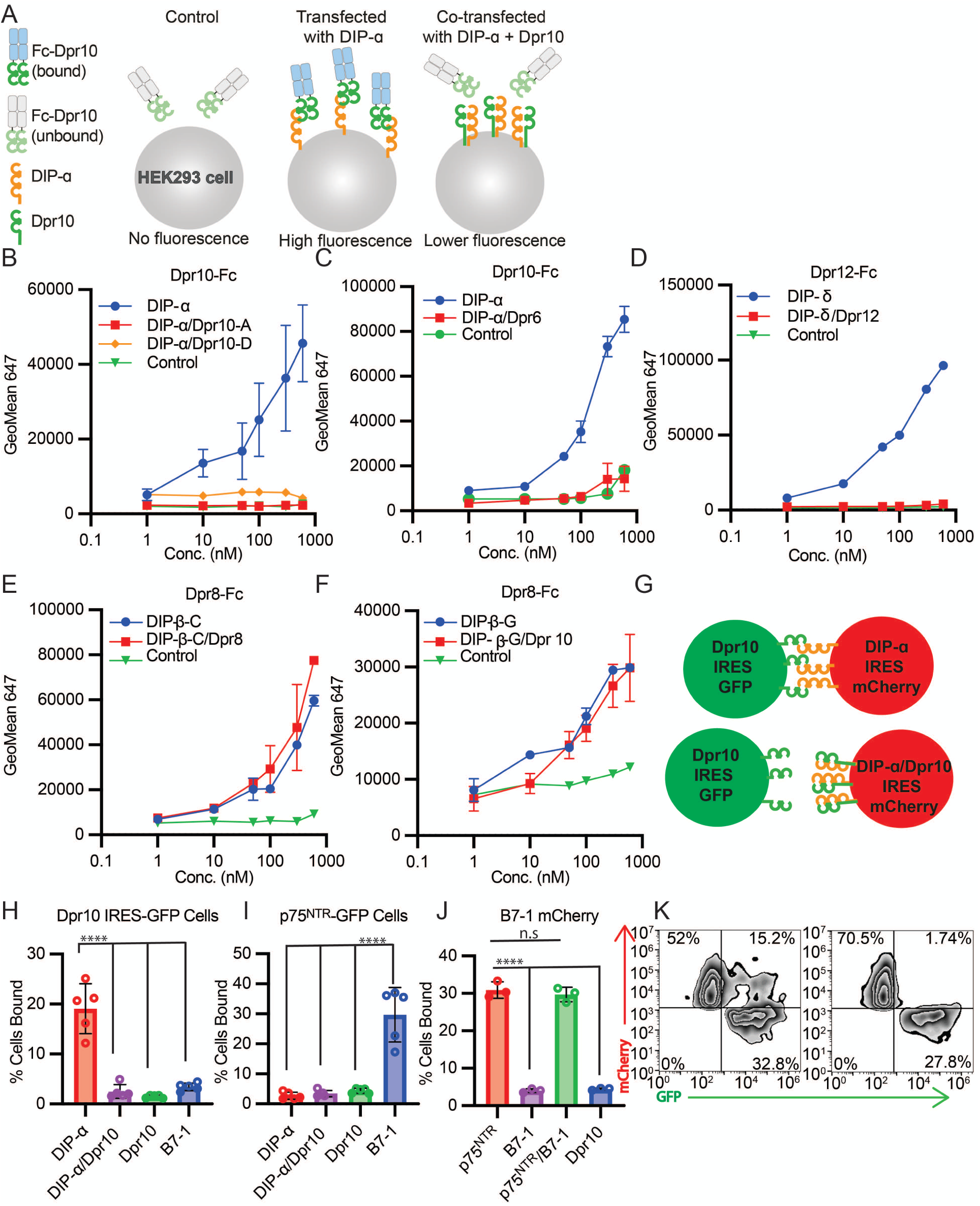
DIP-ɑ interacts in *cis* with Dpr10 and Dpr6. **(A)** Experimental scheme for *cis* inhibition with Fc fusion proteins. Cells are incubated with protein indicated, and then binding is detected with an anti-Fc Alexa 647 labeled antibody using flow cytometry. Data is analyzed by plotting geometric mean of 647 signal of cells positive for both GFP and mCherry. **(B)** Titration of Dpr10-Fc from 1-600 nM onto HEK293F cells expressing DIP-ɑ IRES mCherry, co-expressing DIP-ɑ IRES mCherry and Dpr10-A, co-expressing DIP-ɑ IRES mCherry and Dpr10-D, or control cells (transfected with GFP). **(C)** Same as **(B)** but with Dpr6 co-transfected. **(D)** Dpr12-Fc was titrated from 1-600 nM onto HEK293F cells expressing DIP-δ IRES mCherry, co-expressing DIP-δ IRES mCherry and Dpr12, or control (transfected with GFP) cells. **(E)** Dpr8-Fc was titrated from 1-600 nM onto HEK293F cells expressing DIP-β-C IRES mCherry, co-expressing DIP-β-C IRES mCherry and Dpr8, or control cells transfected with GFP. **(F)** Same as **(E)** but with DIP-β-G IRES mCherry co-transfected with Dpr10-A. **(G)** Experimental scheme for *cis* inhibition using cell aggregation. Cells expressing interacting/noninteracting pairs and different FPs were incubated together and aggregation were assessed by flow cytometry. Data is presented as % of FP positive cells that are positive for both GFP and mCherry. **(H)** Cells expressing Dpr10 IRES GFP or **(I)** p75^NTR^-GFP were queried against cells expressing DIP-ɑ, Dpr10, both DIP-ɑ and Dpr10, or B7-1, all tagged with mCherry. **(J)** B7-1-mCherry was queried against cells expressing p75^NTR^-GFP, p75^NTR^-GFP and B7-1-GFP, or Dpr10 IRES GFP. Unlike in the DIP-ɑ::Dpr10 system, no difference in binding was observed to cells expressing both p75^NTR^-GFP and B7-1-GFP.**(K)** Representative Zebra-plot demonstrating binding differences of DIP-ɑ IRES mCherry cells with (left) Dpr10 IRES GFP expressing cells and (right) DIP-ɑ/Dpr10 IRES mCherry cells. All data was analyzed using one way ANOVA with multiple comparisons, n = 5, **** = p<.0001.

The phenomenon of *cis* inhibition was not observed for all pairs of DIPs and Dprs. For example, binding of Dpr8-Fc was detected to cells expressing DIP-β-C or co-expressing DIP-β-C and Dpr8 (**Figure 2E**). Similarly, Dpr8-Fc was able to bind to cells expressing DIP-β-G or cells co-expressing DIP-β-G and Dpr10-A (**Figure 2F**). The lack of *cis* inhibition in these cases is notable because both DIP-β-G::Dpr10 and DIP-β::Dpr8 are high affinity interactions. We show below that the ability to inhibit in *cis* depends on the length of the linker that separates the most proximal Ig domain from the cell membrane.

### Cell-cell flow cytometry experiments also show *cis* inhibition of *trans* binding

In light of the above results, we next tested whether *cis* inhibition can be detected in our flow cytometry assay (as shown in **Figure 1, schematized in Figure 2G**). These experiments were performed by mixing cells that express Dpr10-IRES-GFP with cells transfected with either DIP-α-IRES-mCherry, Dpr10-IRES-mCherry, or both. Additionally, B7-1 and p75^NTR^, which are known to form a complex, were included as positive and negative controls for cell aggregation. While we detected cell aggregation between cells expressing Dpr10 and cells expressing DIP-α, co-expression of both DIP-α and Dpr10 ablated aggregation with Dpr10 expressing cells (**Figure 2H-K**). In contrast, p75 expressing cells aggregated with cells expressing its cognate ligand B7-1 and also cells expressing both p75 and B7-1 (**Figure 2I-J**). Consistent with our results for cell-protein binding, these studies suggest that DIP-α interactions with Dpr10 in *cis* can inhibit *trans* interactions between DIP-α and Dpr10, and that not all adhesion molecules exhibit *cis* inhibition.

### *Cis* inhibition exploits the *trans* interface

To determine whether DIP-α::Dpr10 *cis* interactions occur via the same interface used by *trans* interactions **(Figure 3A)**, we examined whether a Dpr10 protein harboring the Y103D mutation, previously shown to disrupt the *trans* interface, can inhibit when expressed in *cis*. We titrated Dpr10-Fc onto cells co-expressing DIP-α and Dpr10-Y103D, as well as cells co-expressing DIP-α and Dpr12 (to control for possible changes in DIP-α expression). Dpr10-Fc binding was detected to cells expressing both co-transfected pairs, but not to cells co-expressing of DIP-α and WT Dpr10 (**Figure 3B**), suggesting that *cis* inhibition requires the same interface used for *trans* binding between these proteins.

**Figure 3:**
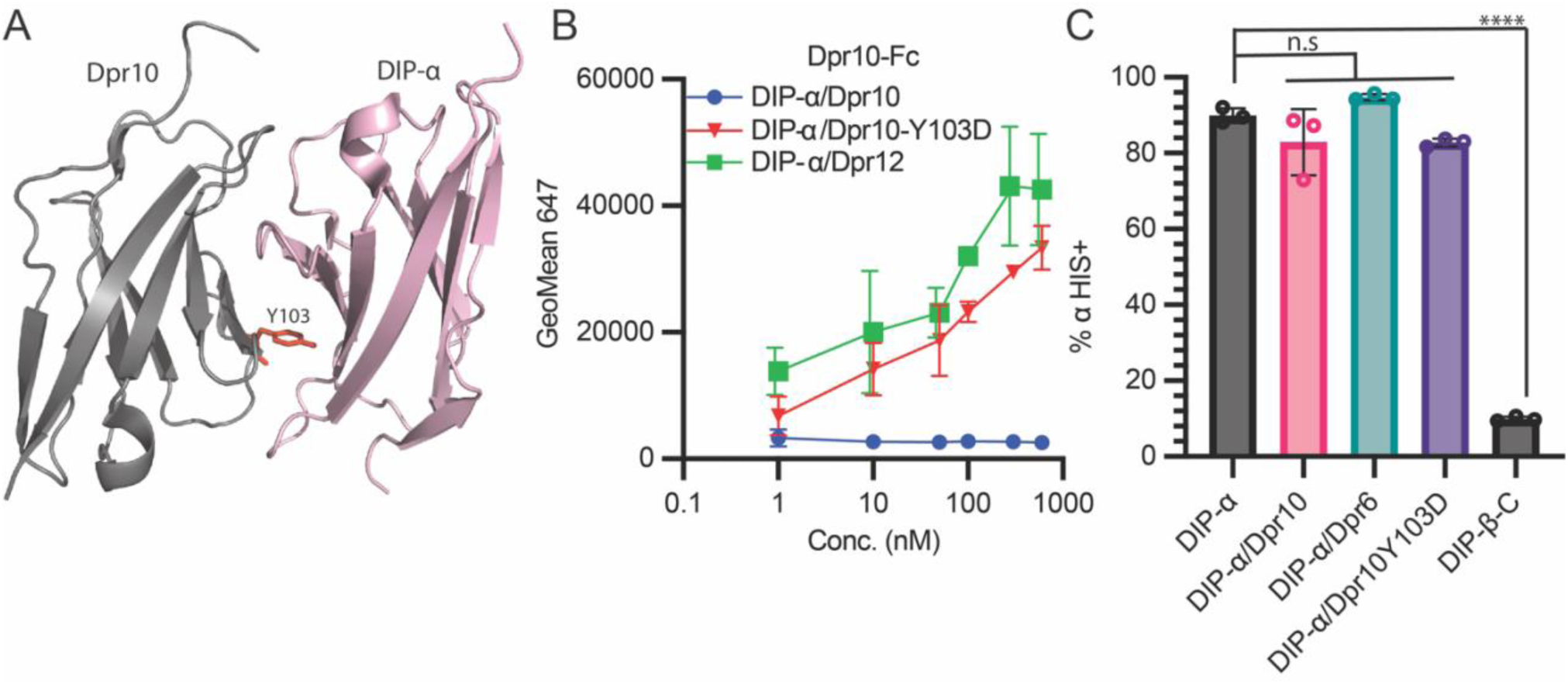
DIP-ɑ induces *cis* inhibtion with Dpr10 and Dpr6 through the *trans* interface. **(A)** Ribbon representation of DIP-ɑ bound to Dpr10 (PDBID: 6NRQ). Mutation of Dpr10 Y103 residue (shown in red sticks) into aspartate disrupts the *trans* interface. **(B)** Dpr10-Fc was titrated from 1-600 nM onto HEK293F cells co-expressing DIP-ɑ IRES mCherry and Dpr10, co-expressing DIP-ɑ IRES mCherry and Dpr12, or co-expressing DIP-ɑ IRES mCherry and Dpr10-Y103D. **(C)** Surface staining of DIP-ɑ transfected alone, or co-transfected with Dpr10, Dpr6, or Dpr10-Y103D could be visualized with an anti HIS antibody (which interacts with poly HIS region of linker domain). DIP-β is included as a negative control. (All data was analyzed using one way ANOVA with multiple comparisons, n = 5, **** = p<.0001).

### *Cis* inhibition occurs on the cell surface

The above results do not differentiate if *cis* inhibition is a consequence of binding at the cell surface, or if binding occurs during trafficking in the ER/Golgi, possibly preventing cell surface localization. Ideally, addressing this question would require direct labeling of DIP-α. However, there are no readily available reagents that bind to DIP-α, and inserting a tag into the protein could alter its biophysical properties. Fortuitously, the C-terminal region of DIP-α contains the amino acid sequence HHHHHHH N-terminal to the predicted GPI anchor site, which makes it possible to detect cell surface DIP-α with a fluorescently labeled anti-6xHis antibody. Using this approach, we observed the presence of cell surface DIP-α on live cells expressing DIP-α, or co-expressing DIP-α and Dpr10, DIP-α and Dpr6, or DIP-α and Dpr10-Y103D. DIP-β was included as a negative control. While DIP-β displayed no antibody binding (as expected due to the lack of an HHHHHHH sequence), there were no differences in cell surface levels of DIP-α under any of the other conditions (**Figure 3C**). These results suggest that co-expression of Dpr10 does not prevent the trafficking and cell surface localization of DIP-α, and suggests that *cis* inhibition occurs on the cell surface.

### Long linker lengths are needed for *cis* binding between cognate DIPs and Dprs

A striking feature of some DIPs is the presence of long extracellular stretches of amino acids C-terminal to the Ig domains. Notably, secondary structure prediction methods suggest that these regions are largely disordered. Recent experimental evidence indicates that *Drosophila* DIPs and Dprs are GPI-anchored to the cell membrane^26^. As a consequence, the effective linker length connecting structured domains to the membrane corresponds to the number of residues between the C-terminus of the membrane proximal Ig domain, Ig3 for DIPs and Ig2 for Dprs, and the ω-site.

Prior to testing if linker length is relevant to *cis* inhibition, we first identified the most plausible location of the GPI site that anchors these proteins to the cell membrane. To identify ω-sites in proteins with ambiguous GPI signatures in their C-terminal regions, we employed an adapted protocol for GPI site prediction (see Methods; **Supplementary Figures S5, S6, Supplementary Data File 2).** The predicted ω-sites, along with the protein linker lengths to the membrane, are detailed in **Figure 4A** (based on data presented in **Supplementary Figure S6**). In *Drosophila melanogaster*, DIP-α is predicted to be GPI-anchored at S501, which results in a linker length of 161 amino acids between its Ig3 domain and the membrane. Conversely, DIP-β’s GPI anchor prediction is located at S458 with a notably shorter linker of 52 amino acids. DIP-δ possesses a predicted linker length of 112 amino acids, intermediate relative to DIP-α and DIP-β. In contrast, Dprs 6, 8, 10 and 12 are predicted to have relatively short linker regions (**Figure 4A**). Sequence alignments of C-terminal regions of DIPs and Dprs for multiple *Drosophila* species (see Methods for details) reveal that the ω-sites and linker lengths are well-conserved (**Figure 4A** and **Supplementary Figure S6B**).

**Figure 4:**
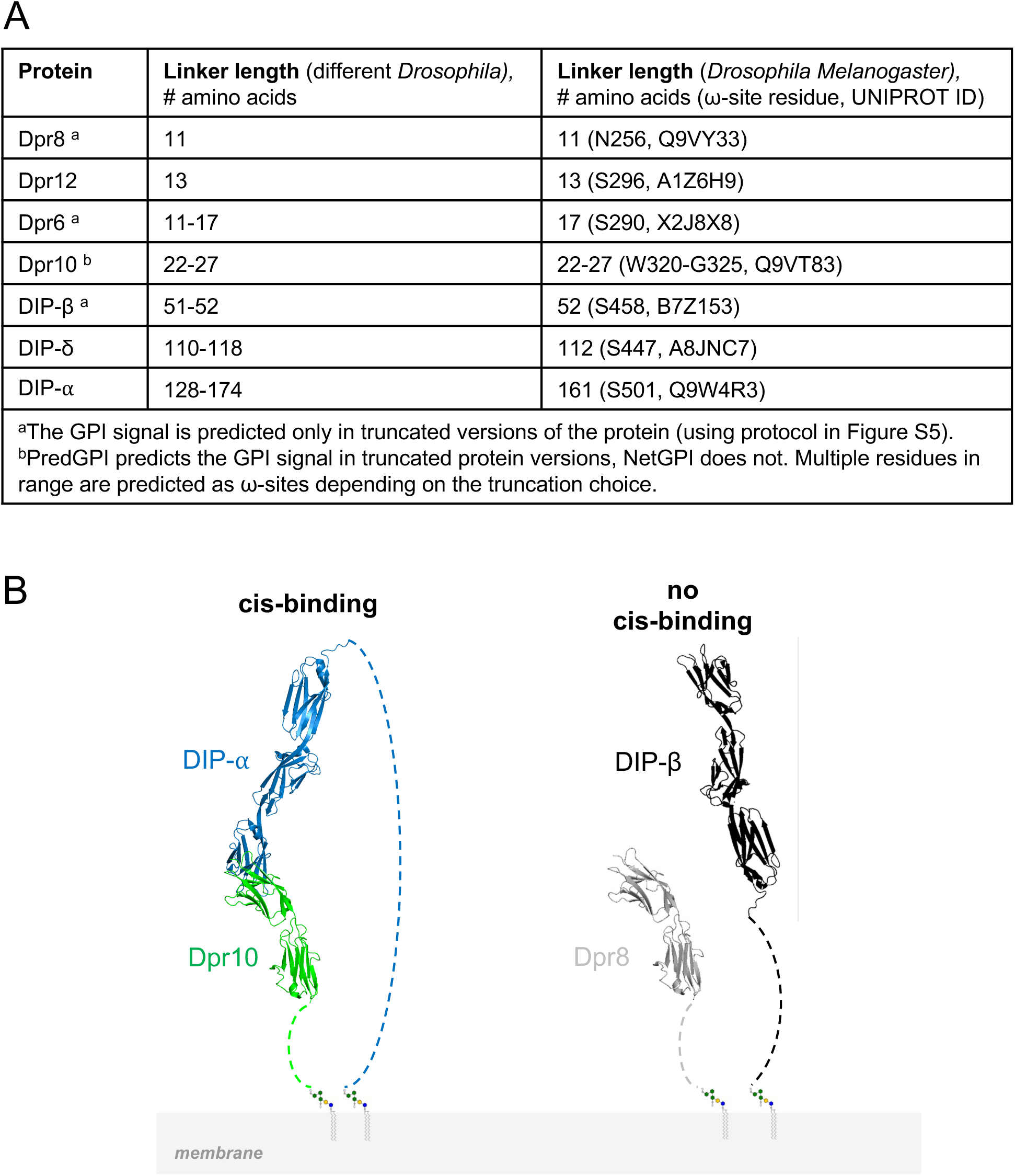
Linker length conservation and requirements in *Drosophila.* **(A)** Results of sequence analysis showing that linker lengths are conserved within different *Drosophila* species. **(B)** The left panel indicates that the linker lengh in DIP-〈 is long enough to span the five Ig domains of DIP-〈::Dpr10 complex. The right panel shows why the short linker in DIP-® is not able to accommodate *cis* binding. Extracellular domains of DIPs and Dprs are shown in ribbon representation. C-terminal tails connecting membrane proximal domains to the GPI anchor as dashed lines.

To confirm the identification of ω sites, we generated two mutations for both DIP-α and DIP-β. The first (del-GPI) truncates both proteins from the start of the predicted GPI anchor ω site to the C-terminus. The second set of mutants disrupts the hydrophobic region of the GPI anchor signal motif for both proteins by mutating Leu-Leu to Glu-Glu (**Figure 5A-B**). Both constructs were transfected into HEK293 cells and expressed mCherry via the IRES signal. 600 nM of Dpr10-Fc or Dpr8-Fc was incubated with either wildtype or mutated DIP-α and DIP-β, and binding was assessed via a 647-labeled antibody and detected via flow cytometry (**Figure 5A-B**). While binding was detected to the wildtype proteins, neither mutant bound to their cognate Dpr for DIP-α and DIP-β. This suggests that these mutations perturb cell surface localization, and eliminate their predicted cell surface linkage.

**Figure 5:**
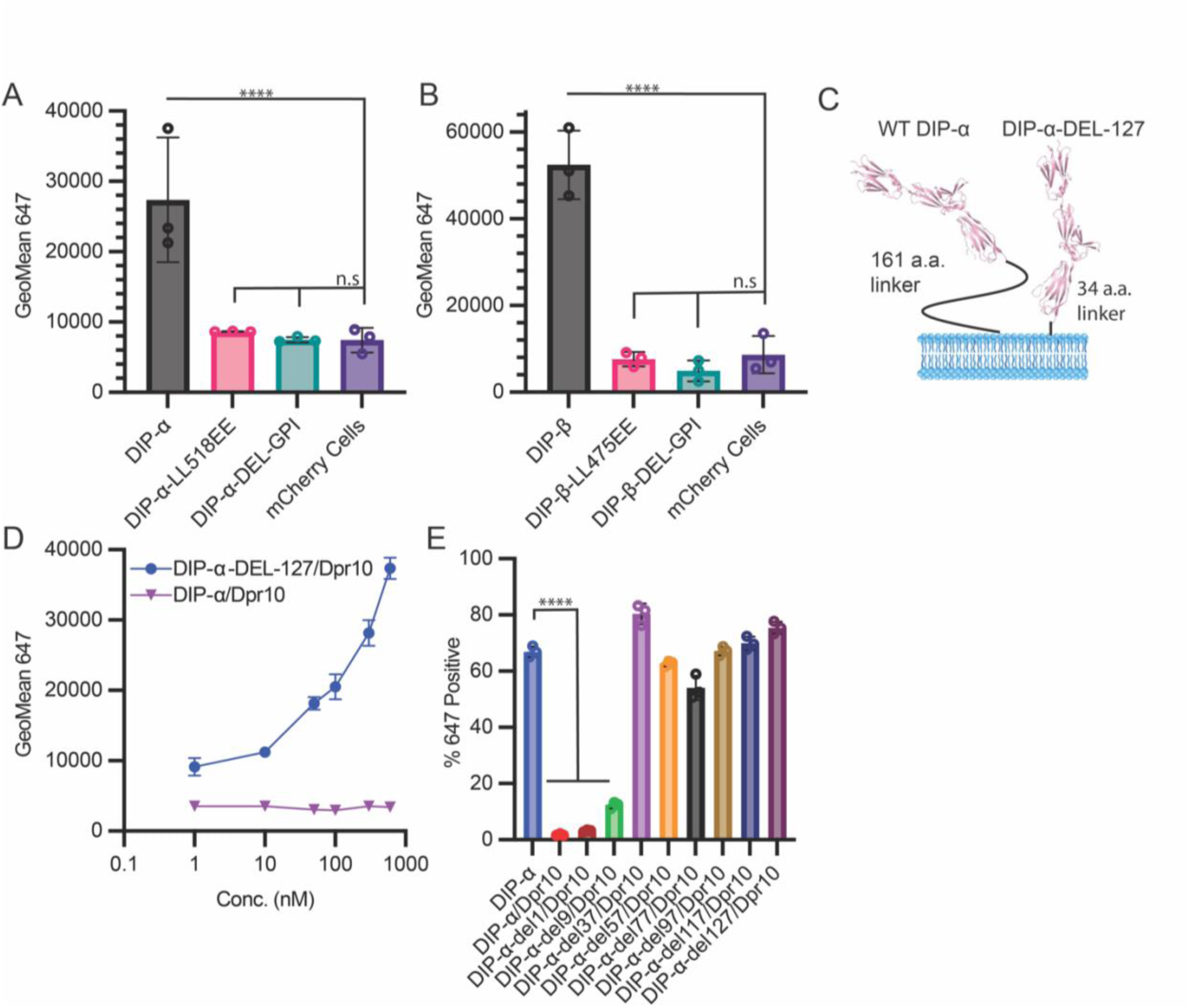
Long DIP-ɑ Linker length is requireed for *cis* binding. **(A)** Dpr10-Fc binding to WT DIP**-**α and DIP**-**α hydrophobic GPI (+) region mutant (LL518EE), and truncation of GPI site mutant (DelGPI). Binding, assessed as described in Figure 2, was not detected to either mutant. **(B)** Dpr8-Fc binding to WT DIP**-**β and DIP**-**β hydrophobic GPI tail mutant (LL475EE), and truncationoo of GPI site mutant (DelGPI). Binding was not detected to either mutant. **(C)** Cartoon of a DIP-ɑ linker mutant that has its linker region truncated to 34 amino acids, compared to wildtype. **(D)** Titration of Dpr10-Fc onto cells expressing either WT DIP-ɑ-L34/Dpr10 or DIP-ɑ/Dpr10. Binding was not detected to DIP-ɑ/Dpr10, but was detected to DIP-ɑ-L34/Dpr10, conducted as in Figure 2. **(E)** A series of truncations of the DIP-ɑ linker region (resulting in linker lengths between 34 to 152 amino acids) were made and assessed for *cis* inhibition by incubating with 300 nM Dpr10-Fc. Dpr10-Fc binding was observed with all co-transfections, except WT-DIP-ɑ/Dpr10 and DIP-ɑ-L152/Dpr10 (All data was analyzed using one way ANOVA with multiple comparisons, n = 3, **** = p<.0001).

How might linker length affect binding in *cis*? We hypothesize that since *cis* and *trans* binding use the same interface, a long linker may be required to enable *cis* binding in a *trans*-like orientation. The difference in linker length could explain differences between DIP-α/δ and DIP-β in their ability to inhibit in *cis*. As illustrated in **Figure 4B**, the linker would have to be long enough to span 5 Ig domains which, in the crystal structure (PDBID: 6EG0)^9^, corresponds to about 165 Å. While the ensemble of conformations that are adopted by these linker regions is unknown, it is possible that longer linkers facilitate the formation of *cis* interactions. To test this hypothesis, a series of truncations were designed in which portions of the C-terminal linker of DIP-α were replaced by the three amino acids G-G-S, resulting in linkers of lengths 152, 124, 104, 84, 64, 44, or 34 amino acids (termed DIP-α-L152, DIP-α-L124, etc.). All truncations removed regions between Ig3 and the GPI anchor, to ensure that GPI anchorage and Ig domains remained intact. When co-transfected with Dpr10, all mutants maintained strong binding to Dpr10-Fc except for the 160 and 152-length linkers, indicating that shortened linkers prevent *cis* inhibition (**Figure 5C-E**). Of note, DIP-α-L152, with a 124 amino acid long linker, cannot inhibit binding to Dpr10-Fc while WT DIP-δ, with a 112 aa linker, inhibits binding to Dpr12 (**Figure 2D**), suggesting that linker length may not be the only parameter that impacts *cis* inhibition.

### *Cis* inhibition in the developing mushroom body

As an initial test to determine if *cis* inhibition between DIPs and Dprs can occur *in vivo,* we first turned to the *Drosophila* mushroom body (MB), where we recently established a role for Dpr12::DIP-δ *trans*-neuronal interactions during development^14^. The MB is comprised of three types of sequentially-born intrinsic neurons - γ, ɑ’/β’ and ɑ/β, collectively known as Kenyon cells (KCs). The first-born γ-KCs undergo stereotypic remodeling during metamorphosis, which includes pruning of the larval axonal lobes, followed by regrowth of an adult-specific lobe^27^. Extrinsic neurons of the MB circuit, including MB output neurons (MBONs) and modulatory dopaminergic neurons (DANs) of two main clusters, innervate KC axons at discrete locations, thus defining distinct sub-axonal zones along the MB lobes. In the adult γ-lobe they define 5 zones termed γ1-γ5 ^28,29^ (**Figure 6A**). We previously discovered that formation of the adult γ4/5 zones is mediated by *trans*-neuronal interactions between Dpr12 in γ-KCs and DIP-δ in DANs of the PAM cluster (PAM-DANs). Loss of either member of the cognate pair results in termination of γ-axon regrowth at the γ3-γ4 border, and thus failure to form the γ4/5 zones^14^ (see model in **Figure 6A** and representative images in **B-C**).

**Figure 6:**
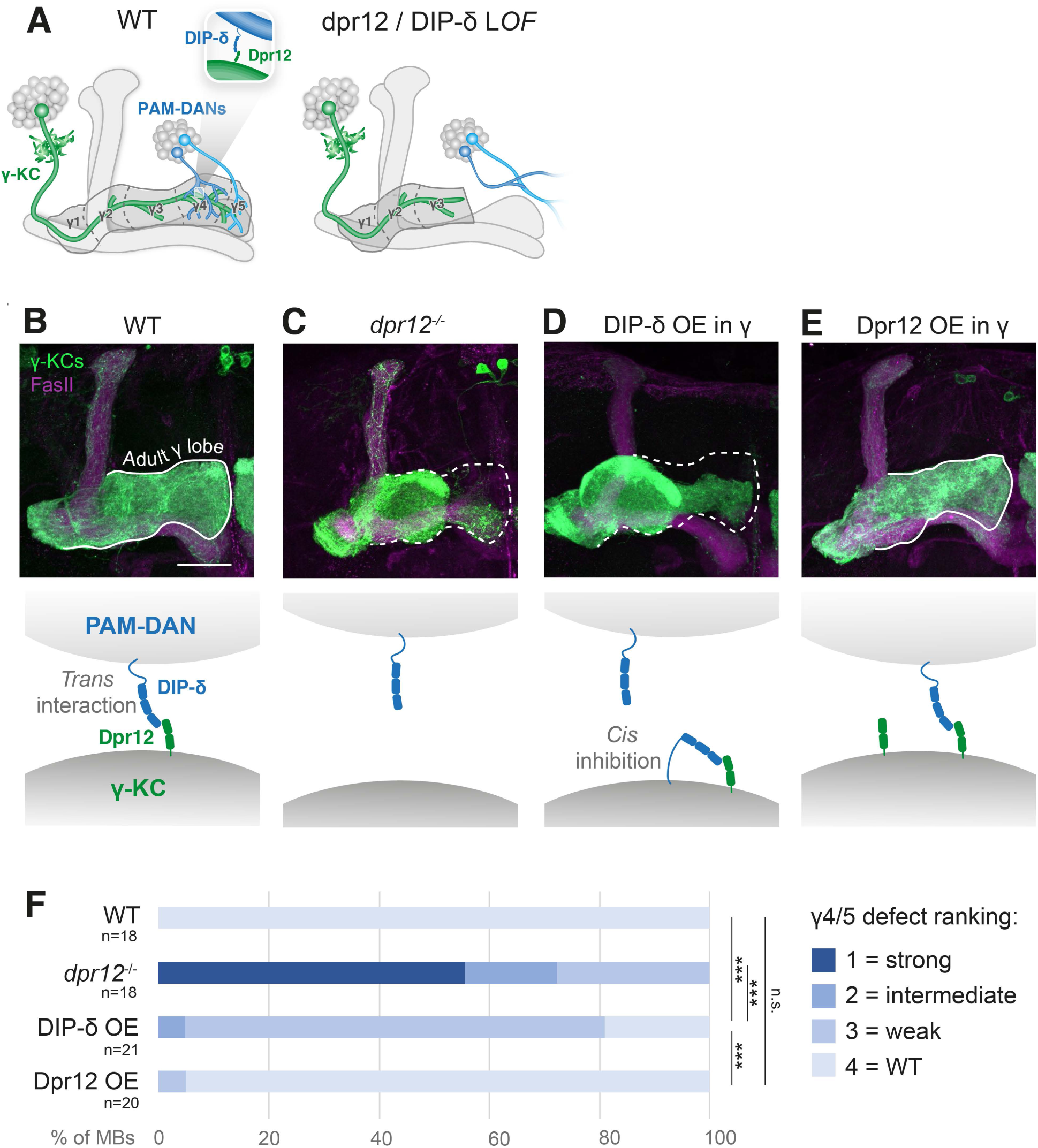
DIP-δ mis-expression in mushroom body γ-KCs mimics Dpr12 loss-of-function defective zone formation. **(A)** Schematic illustration of the adult mushroom body circuit in WT flies vs. Dpr12/DIP-δ mutants. γ-KCs, γ-Kenyon cells (green); PAM-DANs, dopaminergic neurons of the PAM-cluster (blue). Adapted from Bornstein et al. 2021.**(B-E)** Confocal z-projections of adult MBs in which the γ-specific R71G10-Gal4 drives expression of membrane-bound GFP (green), in WT flies (B); dpr12 homozygous mutants (C); or upon R71G10-Gal4-driven overexpression of either DIP-δ (D) or Dpr12 (E). The adult γ-lobe is outlined by a solid white line when regrowth is normal (B, E). A dashed line indicates the putative adult γ-lobe in cases in which regrowth is (at least partially) arrested at the γ3-γ4 border (C, D). Magenta is FasII staining, which strongly labels ɑ/β-KCs and weakly labels γ-KCs. Scale bar is 30µm. The schemes below illustrate the hypothesized model in each scenario.**(F)** Quantification of the γ4/5 defect phenotypes in B-E, ranging from strong defect (light blue) to WT-like morphology (dark blue). *** is p<0.001.

To test whether *cis* inhibition can occur in this system, we mis-expressed DIP-δ in γ-KCs (which endogenously express Dpr12 but not DIP-δ). This manipulation indeed induced an axon extension defect resulting in malformation of the γ4/5 zones, which, although milder, mimics the phenotype of *dpr12* mutants (**Figure 6B-D**, quantified in **F**). A plausible explanation for this effect is that the mis-expressed DIP-δ binds Dpr12 in *cis* within the γ-axon membranes, thereby ‘sequestering’ Dpr12 and reducing its availability to *trans*-interact with DIP-δ in PAM-DANs. Once *trans*-neuronal interactions between γ-KCs and PAM-DANs are compromised, the γ4/5 zones cannot properly form. Notably, this effect seems to be specific to DIP-δ, since, for example, overexpressing Dpr12 in γ-KCs did not affect zone formation and results in WT-like morphology (**Figure 6B** vs. **E**, quantified in **F**).

### Dpr6 and Dpr10 interact with DIP-α in *cis* to regulate leg motor neuron terminal branch morphology in *Drosophila*

To further test for a potential role of *cis* inhibition *in vivo*, we focused on three adult leg MNs, referred to here as the αMNs. The αMNs require a *trans* interaction between DIP-α, expressed in the αMNs, and Dpr10, expressed in leg muscles, for their terminal branching **(Figure 7A-C)** ^12^. Using MiMIC Gal4 insertions to follow the expression of these genes, we find that *dpr6* and *dpr10* are expressed broadly in many leg MNs **(Supplementary Figure S7A-B).** To confirm that these include the DIP-α-expressing αMNs, we analyzed a single cell (sc) RNA-seq dataset of ∼29 leg MNs that includes the αMNs at four timepoints: late 3rd instar, 20 hours after pupal formation (hrs APF), 45 hrs APF, and adult. After identifying the three αMNs (targeting the long tendon <u>m</u>uscle in the <u>fe</u>mur, αFe-ltm; the long tendon <u>m</u>uscle in the <u>ti</u>bia, αTi-ltm; and the tarsal <u>d</u>epressor <u>m</u>uscle in the <u>ti</u>bia, αTi-tadm; **Figure 7B,C**), we confirmed that they co-express *DIP-α, dpr6* and *dpr10* (**Figure 7D-G**). The expression of *DIP-α* and *dpr6* peaks at ∼45 hrs APF, while the expression of *dpr10* peaks at 20 hrs APF in all three αMNs. The co-expression of cognate DIP and Dpr pairs raises the possibility that *cis* inhibition may be playing a functional role in these neurons.

**Figure 7:**
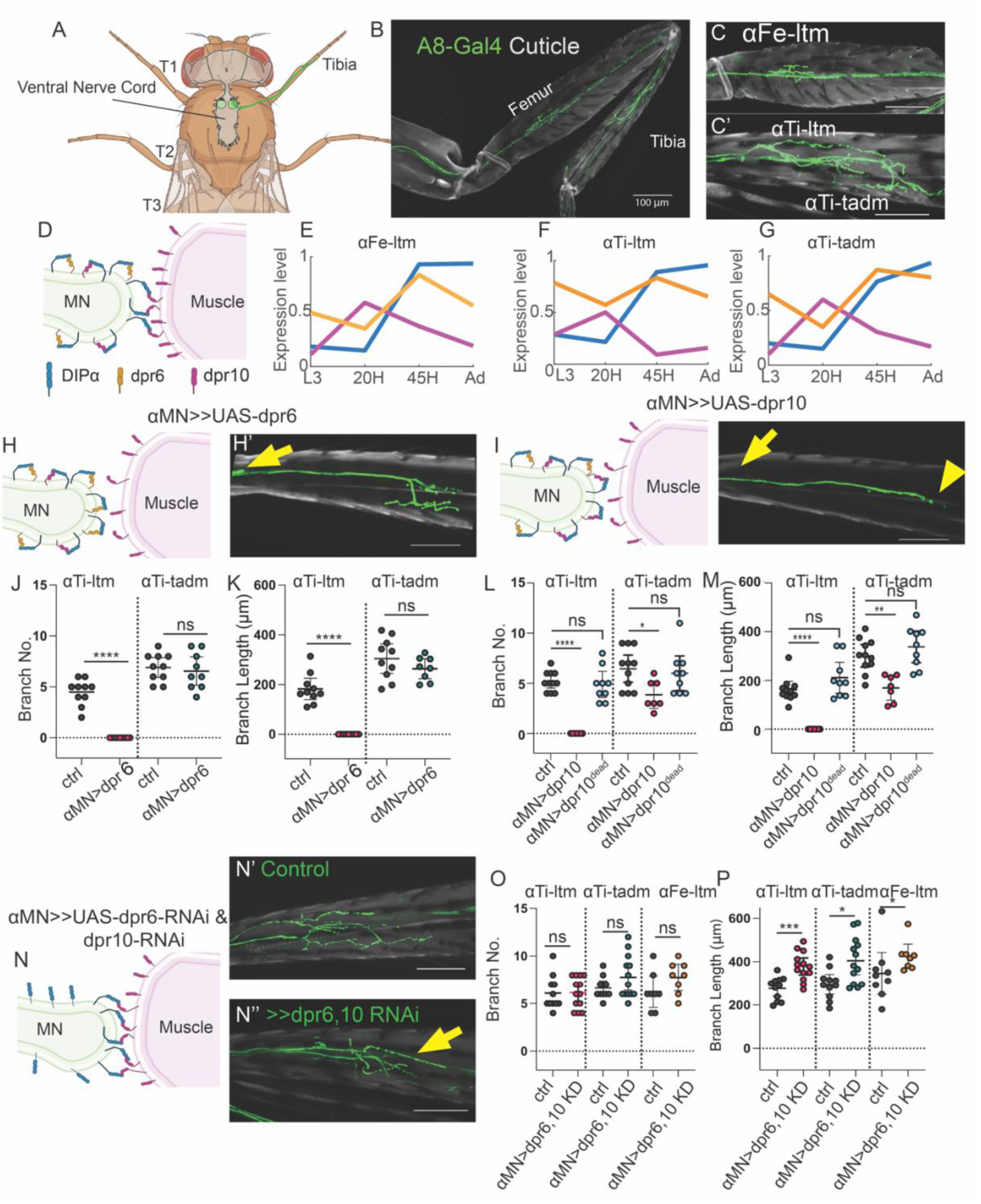
Coexpression of Dpr6, Dpr10, and DIP-α in MNs inhibits Dip-α–Dpr10 interactions in *trans* to regulate leg motor neuron morphology. **(A)** Schematic illustration of *Drosophila melanogaster* central nervous system and leg MNs. **(B)** A8-Gal4>>20XUAS6XGFP expressed in *DIP-α* expressing neurons (αMNs). **(C)** αMNs in Femur, αFe-ltm. (scale bar= 100µm). **(C’)** αMNs in Tibia, αTI-ltm and αTI-tadm. (scale bar= 100µm). **(D)** Model of *cis* regulation of *dpr6* and *dpr10* and *DIP-α*. **(E-G)** Timecourse of relative levels from scRNAseq measurements for *dpr6*, *dpr10* and *DIP-α* in αFe-ltm **(E)**, αTi-ltm **(F)** and αTi-tadm **(G)** color coded according to D. (H) *cis* inhibition of *dpr6* and *DIP-α.* (H’) Representative image of *DIP-α-T2A-Gal4>>20XUAS6XGFP, UAS-dpr6* tibia. (scale bar= 100µm). Yellow arrow shows absence of αTI-ltm terminal branches. (I) *cis* inhibition of *dpr10* and *DIP-α* (I’) Representative image of *DIP-α-T2A-Gal4>>20XUAS6XGFP, UAS-dpr10* tibia. Yellow arrow shows absence of αTI-ltm terminal branches. Yellow arrowhead absence of αTI-tadm terminal branches. (scale bar= 100µm). **(J)** Quantification of the number of αTi-ltm and αTi-tadm terminal branches.Ctrl, *DIP-α-T2A-Gal4>>20XUAS6XGFP.* αTi-ltm n= 10. αTi-tadm n=10 *αMN>dpr6, DIP-α-T2A-Gal4>>20XUAS6XGFP,UAS-dpr6.* αTi-ltm n=9. αTi-tadm n=9. **(K)** Quantification of the sum of αTi-ltm and αTi-tadm branch length.Ctrl, *DIP-α-T2A-Gal4>>20XUAS6XGFP.* αTi-ltm n= 10. αTi-tadm n=10. *αMN>dpr6, DIP-α-T2A-Gal4>>20XUAS6XGFP,UAS-dpr6.* αTi-ltm n=9. αTi-tadm n=9. **(L)** Quantification of the number of αTi-ltm and αTi-tadm terminal branches.Ctrl, *DIP-α-T2A-Gal4>>20XUAS6XGFP.* αTi-ltm n=11. αTi-tadm n=11. *DIP-α-T2A-Gal4>>20XUAS6XGFP,UAS-dpr10.* αTi-ltm n=7. αTi-tadm n=7. *DIP-α-T2A-Gal4>>20XUAS6XGFP,UAS-dpr10Y103D.*αTi-ltm n=9. αTi-tadm n=9. **(M)** Quantification of the sum of αTi-ltm and αTi-tadm branch length. Ctrl, *DIP-α-T2A-Gal4>>20XUAS6XGFP.* αTi-ltm n=11. αTi-tadm n=11. *DIP-α-T2A-Gal4>>20XUAS6XGFP,UAS-dpr10.*αTi-ltm n=7. αTi-tadm n=7. *DIP-α-T2A-Gal4>>20XUAS6XGFP,UAS-dpr10Y103D.*αTi-ltm n=9. αTi-tadm n=9. **(M)** Quantification of the sum of αTi-ltm and αTi-tadm branch length. **(N)** Double knockdown (KD) of *dpr6* and *dpr10* in MNs (in *cis*). (N’) A8-Gal4>>20XUAS6XGFP control animals. (scale bar= 100µm). **(N’’)** A8-Gal4>>20XUAS6XGFP, UAS-dpr6-RNAi, UAS-dpr10-RNAi. (scale bar= 100µm). Yellow arrow indicates elongated branches. **(O)** Quantification of the number of αFe-ltm, αTi-ltm, αTi-tadm terminal branches. Ctrl, *A8-Gal4>>20XUAS6XGFP.* αTi-ltm n=11. αTi-tadm n=11. αFe-ltm n=9. *αMN>dpr6,10KD, A8-Gal4>>20XUAS6XGFP,UAS-dpr6-RNAi, UAS-dpr10-RNAi*. αTi-ltm n=13. αTi-tadm n=13. αFe-ltm n=10. **(P)** Quantification of the sum of αFe-ltm, αTi-ltm and αTi-tadm branch length. Ctrl, *A8-Gal4>>20XUAS6XGFP.* αTi-ltm n=11. αTi-tadm n=11. αFe-ltm n=8. *αMN>dpr6,10KD A8-Gal4>>20XUAS6XGFP,UAS-dpr6-RNAi, UAS-dpr10-RNAi*αTi-ltm n=13. αTi-tadm n=13. αFe-ltm n=8. For all graphs statistical significance was determined using an unpaired nonparametric two tailed Mann-Whitney test. Error bars represent mean with 95% confidence intervals. ns= no statistical difference. *p<0.05 **p<0.01 ***p<0.001 ****p<0.0001 *T1-Prothoracic segment*, *T2-mesothoracic segment*, *T2-metathoracic segment*, *L3-Late 3rd instar, H-Hours after pupal formation. Ad-Adult Ctrl-Control, MN-motor neuron, KD-knockdown*.

If *cis* inhibition is occurring in these MNs, we reasoned that altering the ratio of *Dpr10:DIP-α* and *Dpr6:DIP-α* in αMNs might affect the ability of their terminal branches to form stable *trans* interactions with *dpr10* on muscles. A Gal4 insertion into *DIP-α* that is expressed from 20 hrs APF onwards in αMNs was used to overexpress either *dpr6* or *dpr10,* and the morphology of two of the three αMNs, αTi-ltm and αTi-tadm, was examined (**Figure 7H-I’**). Overexpressing either *dpr6* or *dpr10* in αMNs inhibited the formation of almost all αTi-ltm terminal branches **(Figure 7J-M).** This phenotype resembles the *DIP-α* and *dpr10* null phenotypes^12^, where αTi-ltm axons still reach their targets but fail to form terminal branches. The effect on αTi-tadm’s terminal branches is weaker, and it is only observed with *dpr10* but not *dpr6* overexpression. The different effects on these two MNs parallel the different penetrance of the terminal branching phenotype of *DIP-α* and *dpr10* null mutants, where both Ti-ltm and Ti-tadm have different sensitivities to loss of DIP-α^12^. We also note that differences in the affinities between Dpr6 and Dpr10 to DIP-α (1.67 µM and 2.06 µM respectively)^9^ and/or in the timing or cell surface distribution of DIP-α could also contribute to the different penetrances. Notably, overexpression of Dpr10^Y103D^ in αMNs, which renders it unable to bind DIP-α, did not cause branching defects, confirming that the binding interface is required for *cis* inhibition and that *cis* inhibition is not simply due to overexpressing of Dpr10 **(Figure 7L-M)**. Further, overexpression of Dpr10 in muscles had no effect on the morphology of these MNs (**Supplementary Figure S7C-D**), consistent with the idea that it is the relative abundance of DIP-α:Dpr10 in MNs that regulates binding in *trans*.

Although the above results demonstrate that the ratio of a DIP and its cognate partner within a neuron can affect interactions in *trans*, they do not address whether this mechanism plays a role during normal development. To answer this question, we performed single and double knockdown experiments of *dpr6* and *dpr10* in αMNs **(Figure 7N-P, Supplementary Figure S7E-F)**. We hypothesized that a reduction of *dpr6* and *dpr10* in MNs might promote DIP-α::Dpr10 *trans* interactions and potentially stabilize the interaction between αMNs and muscles. For these experiments, instead of the Gal4 insertion into the endogenous *DIP-α* locus (which is a hypomorphic allele) we used a previously described enhancer fragment from *DIP-α’s* regulatory region called A8, which is specifically expressed in the three αMNs to drive Gal4^12^. Interestingly, although branch number is not affected, the double knockdown of *dpr6* and *dpr10* resulted in longer terminal branches in all three αMNs **(Figure 7P)**. This phenotype was less severe for single knockdowns (**Supplementary Figure S7E-F**). Along the same lines, we would expect that increasing the concentration of DIP-α in αMNs would also change the balance of DIP-α to favor *trans* over *cis* binding and should have a similar phenotype to the *dpr6* and *dpr10* double knockdown. Consistent with this expectation, overexpression of DIP-α in αMNs resulted in an increase in terminal branch length compared to controls for the αTi-ltm and αFe-ltm MNs (**Supplementary Figure S8B)**.

Finally, we also tested the role of linker length in DIP-α’s ability to be inhibited in *cis.* Based on the experiments carried out in cells (**Figure 5)**, the prediction is that if DIP-α’s linker is truncated, *cis* inhibition would be compromised, leading to more DIP-α available for *trans* interactions. To test this we used CRISPR/Cas9 to generate the same 123 amino acid deletion as in DIP-α-L152 in the endogenous *DIP-α* locus, creating the *DIP-α^short^* allele. This allele is functional because, unlike null alleles, the terminal branches for all three αMNs were intact in *DIP-α^short^* homozygote animals (**Figure 8C**). However, the terminal branch lengths of the αTi-ltm were longer compared to controls (**Figure 8C,D**), a phenotype that is shared by other manipulations that reduce *cis* inhibition, such as the double knockdown of *dpr6* and *dpr10*. The increase in branch length was not observed for the αFe-ltm and αTi-tadm, suggesting that *cis* inhibition may not be fully abolished by the *DIP-α^short^* allele.

**Figure 8.**
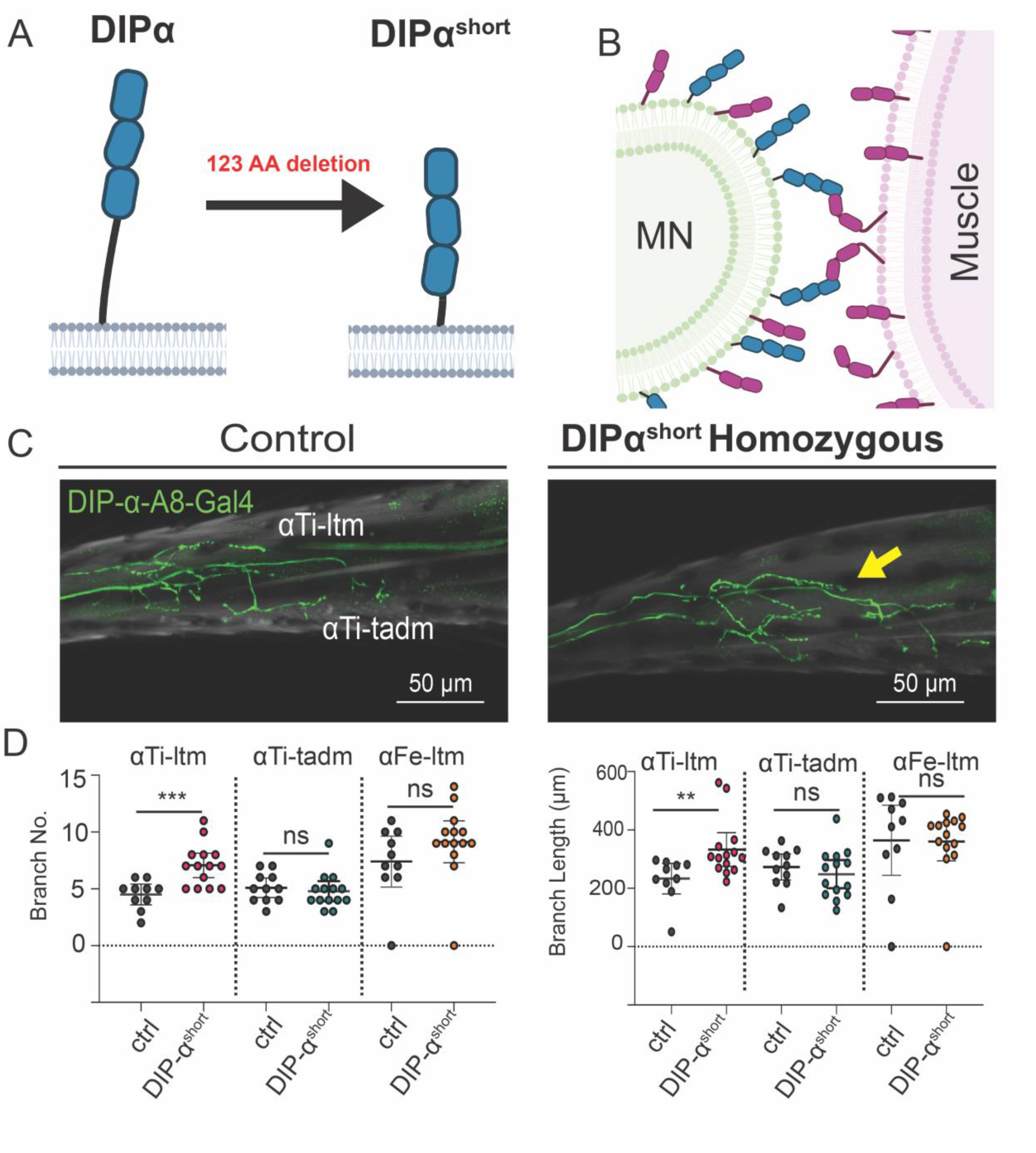
Compromised *cis* inhibition of the *DIP-α^short^* allele in MNs. **A)** Schematic illustration of the truncated DIP-α protein, *DIP-α^short^*. **B)** Model of how *DIP-α^short^* allele can only interact in *trans*. **C)** Representative Images of *DIP-α-A8-Gal4>>20XUAS6XGFP, controls, and DIP-α^short^ homozygous animals, DIP-α-A8-Gal4>>20XUAS6XGFP.*Yellow arrow indicates longer αTi-ltm axon. Scale bar= 50µm **D)** Quantification of the number of αTi-ltm, αTi-tadm and αFe-ltm terminal branches. Ctrl, *DIP-α-A8-Gal4 20XUAS6XGFP.* αTi-ltm n= 11. αTi-tadm n=11 αFe-ltm n=11. *DIP-α^short^ homozygous animals, DIP-α-A8-Gal4>>20XUAS6XGFP..* αTi-ltm n=14. αTi-tadm n=14 αFe-ltm n=14. Quantification of the sum of αTi-ltm, αTi-tadm, and αFe-ltm, branch length. Ctrl, *DIP-α-A8-Gal4 20XUAS6XGFP.* αTi-ltm n= 11. αTi-tadm n=11 αFe-ltm n=11. *DIP-α^short^ homozygous animals, DIP-α-A8-Gal4>>20XUAS6XGFP.* αTi-ltm n=14. αTi-tadm n=14, αFe-ltm n=14. For all graphs statistical significance was determined using an unpaired nonparametric two tailed Mann-Whitney test. Error bars represent mean with 95% confidence intervals. ns= no statistical difference. *p<0.05 **p<0.01 ***p<0.001*. Ctrl-Control, MN-motor neuron*.

In summary, when *cis* inhibition is enhanced, for example by increasing the levels of Dpr10 in αMNs, fewer and/or shorter terminal branches are observed. In contrast, whenever *cis* inhibition is compromised, either by truncating DIP-α’s linker or by increasing the ratio of DIP-α:Dpr10 in MNs, longer terminal branches are observed. Taken together, these consistent changes in αMN terminal branch morphology suggest that *cis* inhibition fine tunes the development of these MNs.

## DISCUSSION

Here, we combine *in vitro* cell assays and *in vivo* genetic manipulations to demonstrate that DIPs and Dprs can interact in *cis* to inhibit *trans* interactions. *Cis* inhibition appears to be a widespread phenomenon among CAMs as it has been observed in several other systems including Notch/Delta^30,31^, ephrins/Ephs^32^, semaphorins/plexins^33^, neurexins/neuroligins^34^, sidekicks^23^, and IgLONs^35–37^, which are the closest vertebrate homologs to the DIPs and Dprs. *Cis* inhibition has also been shown to be important for several developmental processes such as neurogenesis, axon guidance and neural tissue patterning^38,39^. Interestingly, and in contrast to many other examples, DIP::Dpr *cis* and *trans* interactions occur via the same interface, as evident from experiments in which disrupting the well-characterized *trans* interface ablates *cis* inhibition.

We demonstrate *cis* inhibition for two pairs of high affinity DIP::Dpr interactions: Dpr6/10 can inhibit DIP-α, and Dpr12 can inhibit DIP-δ (**Figure 2**). However, we find that high affinity is not sufficient for *cis* inhibition: Isoform G of DIP-β interacts with Dpr10 with a K_D_ of 5.76 µM (**Supplementary Figure S2**) but Dpr10 does not exhibit *cis* inhibition when expressed on the same cell as DIP-β-G **(Figure 2F)**. This implies that some determinants of *cis* inhibition are not localized to the binding interface. Our experiments with truncated linker regions offer a possible mechanism that *cis* inhibition depends on the length of the linker between the C-terminus of the Ig3 domain and the site of the GPI anchor on DIP-α. This linker region is predicted to be largely unstructured and is sufficiently long and flexible to allow for interactions between DIP-α and Dpr10 molecules localized on the same membrane using the same interface as when interacting in *trans*. Notably, there are other examples (including Notch/Delta, sidekicks and IgLONs), that are thought to interact both in *cis* and in *trans* via the same interface. In these cases, either multiple EGF domain repeats (for Notch/Delta) or FN3 repeats (for sidekicks) appear to provide the necessary flexibility. The mammalian IgLONs, which are closely related to DIPs and Dprs^40^, have short linker regions so the mechanism proposed here would not be applicable. However, because all IgLONs have three Ig domains, their orientation on membrane surfaces may permit the formation of a *cis* interaction without the need for a flexible linker^36,37^. Future experiments such as those described here will be required to test this possibility.

While we hypothesize that DIP/Dpr *cis* interactions occur at the cell surface, and that linker length is a key molecular determinant for these interactions, additional mechanisms may also contribute to *cis* inhibition. For example, it is possible that co-expression affects cell surface expression levels, possibly via altering the rate of endocytosis of these receptors. Though our IRES vector based expression strategy has the advantage of expressing full-length, native proteins without tags, it has the caveat of not allowing for direct measurement of cell surface localization. Nevertheless, we show that DIP-α is still trafficked to the cell surface when co-expressed with Dpr10 (**Figure 3C)**, strongly arguing that *cis* inhibition occurs at the cell membrane. The observation that not all DIP-Dpr pairs lead to *cis* inhibition (i.e. DIP-β/Dpr8 and DIP-β/Dpr10), provides further evidence that cell surface levels are not affected by co-expression of interacting pairs. While we point out the difference in linker length between DIP-α, DIP-β, and DIP-δ, *cis* inhibition will need to be tested in all DIP/Dpr pairs to determine the extent of its role in the full DIP/Dpr interactome.

An additional question posed by these experiments is why *cis* interactions appear to almost completely outcompete *trans* interactions despite using the same interface (and thus having the same solution affinity). We suggest that because *cis* interactions occur on the same cell surface, their effective affinity in *cis* is much higher than in *trans* due to a local boost in concentration, thus increasing avidity. We also note that the experiments presented here only investigate end point phenotypes, and do not evaluate the dynamic regulation of *cis* interacting DIPs and Dprs in real time. Such studies using techniques such as single molecule localization microscopy (SMLM)^41^, fluorescence resonance energy transfer (FRET), or proximity ligation assays (PLAs) would potentially be valuable but challenging follow up studies that could directly reveal the dynamic behavior of these receptors in real time.

In addition to mechanistic insights, we also demonstrated that *cis* inhibition impacts fly neurodevelopment. Consistent with our biochemical experiments showing *cis* inhibition of Dpr12 by DIP-δ, we used the well-characterized MB system to show that this inhibition could also be observed *in vivo* (**Figure 6**). Notably, *dpr12* null mutants show a significantly more severe phenotype than that of DIP-δ overexpression. This is consistent with the idea that the *cis* ‘sequestering’ effected by DIP-δ is probably not absolute, and some Dpr12 remains available for *trans* interactions. DIP-δ and Dpr12 are not endogenously co-expressed in MB neurons, and our overexpression experiments do not indicate a role for DIP-δ/Dpr12 *cis* inhibition in normal MB development. Interestingly, other neuronal systems, such as subsets of fruitless (*fruP1*)-expressing neurons^6^, do co-express DIP-δ and Dpr12, implying a potential functional role for *cis* inhibition in these cases.

In the adult fly leg we showed that the relative levels of DIP-α and Dpr6/10 within MNs regulate the morphology of their terminal branches, further supporting the idea that DIP-α and Dpr10/6 interact in *cis in vivo*. We interpret the longer branch lengths observed when *cis* inhibition was compromised, such as when Dpr6 and Dpr10 levels were reduced in MNs, as a consequence of increasing the amount of DIP-α available to bind Dpr10 in muscles, thereby stabilizing the interaction between MN filopodia and muscles. Similarly, replacing the endogenous *DIP-α* allele with one that has a truncated linker, or over-expressing DIP-α in MNs, also resulted in longer terminal branches. Conversely, expressing higher than normal levels of Dpr6/10 in MNs resulted in a decrease in the number of terminal branches. Taken together, these data strongly support the idea that *cis-*expressed Dpr10/6 fine tunes the morphology of MNs that express their cognate partner, DIP-α.

## METHODS

### Generation of DIP/Dpr mCherry and GFP Constructs

Full length constructs of the DIP/Dpr transcript variants (DIP-α isoform A, DIP-β isoforms C and G, DIP-δ isoform E, DIP-ε, and DIP-ζ, Dpr6, Dpr8 isoform B, Dpr9, Dpr10 isoforms A and D, and Dpr12 isoform C) were synthesized by Genscript and Subcloned into psDNA3.1(+) in between the restriction sites NheI and BamHI. All biochemistry experiments were conducted using Dpr10 isoform A unless otherwise noted.The full length genes were then subcloned into the vectors psDNA3.1(+) IRES GFP (NheI and BamHI) (https://www.addgene.org/51406/) and pCI-Neo - IRES mCherry ( EcoRI and XbaI) (https://www.addgene.org/52119/).

### Tissue Culture and Transient Transfection

HEK293 Freestyle suspension cells were cultured in HEK Freestyle Media (Invitrogen, 12338018) grown at 37° C in a humidified shaking platform incubator with 10% CO_2_. For transfection, cells were pelleted at 500xg and resuspended in fresh media. For small-scale (1mL cells at 1×10^6^/mL) transient transfections performed in 24-well non-treated tissue culture plates, 10 μl of 293Fectin (ThermoFisher Cat# 12347019) was added 330 μL of Opti-MEM (ThermoFisher Cat# 31985062), and incubated for 5 min at room temperature. 10 μl of transfection mixture was then added to 1000 ng of DNA, and incubated at room temperature for 30 min, after which it was added to HEK293 Freestyle cells in 24 well plates to 0.5 μg diluted plasmid DNA in a final volume of 100 μL. For co-transfections of DIPs and Dprs, 500 ng of each plasmid was used.

Site directed mutagenesis to generate Dpr10-Y103D, DIP-α-DEL-GPI and DIP-β-LL475EE was performed using high fidelity KOD Hot State polymerase, 2 mM dNTPs and 4mM MgCl_2_ (EMD Millipore, 71086-3). The template used for the mutagenesis included the coding sequence for each DIP/Dpr in the appropriate vector generated above.

### Cell-Cell Binding Experiments

For cell-cell binding experiments, libraries and query ligands were transfected in small scale as described above. One-two days post-transfection, cells were diluted to 1*10^6^ cells/mL in PBS 0.2% BSA, pH 7.4. Binding reactions were setup in 96-well V-bottom plates by mixing equal volumes of challenger (Dpr-IRES GFP expressing cells) and DIP mCherry or DIP/Dpr mCherry expressing cells. After binding, cell-cell conjugates were analyzed by flow cytometry using Novocyte Quanteon (Agilent) or SH800S Cell Sorter (SONY). The percent bound was calculated as the number of double-positive events (GFP and mCherry) divided by the total number of transfected cells. Analysis of flow data was done in FlowJo (FlowJo LLC).

### Surface Plasmon Resonance (SPR) binding experiments

SPR binding assays were performed using a Biacore T100 biosensor equipped with a Series S CM4 sensor chip. DIP-〈, DIP-® (both isoforms) and DIP-⌊ were immobilized over independent flow cells using amine-coupling chemistry in HBS pH 7.4 (10mM HEPES, 150mM NaCl) buffer at 25°C using a flow rate of 20 ⎧L/min. Dextran surfaces were activated for 7 minutes using equal volumes of 0.1M NHS(N-Hydroxysuccinimide) and 0.4M EDC(1-Ethyl-3-(3-dimethylaminopropyl)carbodiimide). Each protein of interest was immobilized at ∼30⎧g/mL in 10 mM sodium acetate, pH 5.5 until the desired immobilization level was achieved. The immobilized surface was blocked using a 4-minute injection of 1.0 M ethanolamine, pH 8.5. Typical immobilization levels ranged between 700-900 RU. To minimize nonspecific binding the reference flow cell was blocked by immobilizing BSA in 10 mM sodium acetate, pH 4.25 for 3 minutes using a similar amine-coupling protocol as described above.

Binding analysis was performed at 25°C in a running buffer of 10 mM Tris-HCl, pH 7.2, 150mM NaCl, 1mM EDTA, 1 mg/mL BSA and 0.01% (v/v) Tween-20. Analytes were prepared in running buffer and using a three-fold dilution series. Dprs 8 and 21 were tested at seven concentrations ranging from 0.012-9 ⎧M. Interactions of Dpr9 with DIP-® isoforms and Dpr10-A with DIP-®-G, DIP-〈 and DIP-⌊ were tested using concentrations ranging from 0.012-27 ⎧M. A more extended concentration range of 0.012-81 ⎧M was used for the remaining interactions to account for their higher K_D_s. Samples were tested in duplicate in order of increasing concentration. A binding cycle consisted of a 30s association phase followed by a 120s dissociation phase, each performed at 50 ⎧L/min, and a buffer wash step of 60s at 100 ⎧L/min. Dpr analytes were replaced by buffer every three binding cycles to double-reference the binding signals by removing systematic noise and instrument drift. The responses between 25 and 29 seconds, at which point the binding reactions achieve equilibrium as observed by the flat binding responses, were plotted against the concentration of analyte. The data was fit to 1:1 interaction model and the K_D_ was calculated as the analyte concentration that would yield 0.5 R_max 42_. The data was processed using Scrubber 2.0 (BioLogic Software).

#### Purification of Recombinant Fc-fusion protein

To clone Dpr-Fc-fusion protein, the first two Ig domains (Dpr10: residues 35 – 236, Dpr8: residues 40 – 244, Dpr12: residues 71 - 285) DNA encoding full length wild type proteins were sub-cloned into a vector containing a C-terminal hexa his-tagged Fc domain (rat IgG1-His6). For Dpr10, the expression constructs as well as mIgG2a and hIgG1 isotype control constructs were transiently expressed in 50 mL of ExpiHEK 293 suspension cells and transfected according to manufacturer guidelines. Seven days post transfection, the media was harvested, 50 mM MES was added to adjust to pH 6.5 and 100 mM Arg-Cl (pH 6.5) was added to enhance solubility. Fc-fusions were subsequently purified by Ni2+His60 chromatography (GE) using a batch binding method (3 mL resin bed volume) followed by gravity flow over a column. The Ni^2+^His60 resin was washed with 3 column volumes of wash buffer (50 mM MES pH 6.5, 100 mM Arg-Cl, 5 mM imidazole, 150 mM NaCl, 10% Glycerol) and the bound protein eluted with 5 mL the same buffer containing 500 mM imidazole. Nickel column elutes were concentrated and further purified by gel filtration on an S200 Sephadex column (MilliporeSigma, GE29321905) equilibrated with 50 mM MES pH 6.5, 100 mM Arg-Cl, 150 mM NaCl, 10% Glycerol. All recombinant proteins were used within one week of purification or were frozen at -80 C and only thawed one time. Frozen aliquots of protein were utilized but routinely checked for potential aggregation by analytical size chromatography). Protein for SPR experiments was produced as previously described^9^.

For DIP-β-G, the sequence below was used:

“FEPDFVIPLENVTIAQGRDATFTCVVNNLGGHRVAWIKADAKAILAIHEHVITNNDRLSVQHNDYNTWTLNIRGVKMEDAGKYMCQVNTDPMKMQTATLEVVIPPDIINEETSGDMMVPEGGSAKLVCRARGHPKPKITWRREDGREIIARNGSHQKTKAQSVEGEMLTLSKITRSEMGAYMCIASNGVPPTVSKRMKLQVHFHPLVQVPNQLVGAPVLTDVTLICNVEASPKAINYWQRENGEMIIAGDRYALTEKENNMYAIEMILHIKRLQSSDFGGYKCISKNSIGDTEGTIRLYEMEHHHHHH”.

For Dpr10-A, the sequence below was used:

“WNEPYFDLTMPRNITSLVGKSAYLGCRVKHLGNKTVAWIRHRDLHILTVGTYTYTTDQRFQTSYHRDIDEWTLQIKWAQQRDAGVYECQISTQPVRSYSVNLNIVDLIDAETSDIMQQYYNDDAFYIAENRVYQSSNDEFAGMFGPIQTVAVPTATILGGPDLYVDKGSTINLTCIIKFSPEPPTHIFWYHQDKVLSEETSGGRLKFKTIKSEETKSILLIYDADLLHSGKYSCYPSNTEIASIRVHVLQGEHHHHHH”.

All other proteins were previously expressed in the above reference.

### Fc-fusion protein Cell titration experiments

Flow cytometry titration assays were performed with Dpr10, Dpr8 and Dpr12 Fc fusion proteins purified as described above. Cells were transfected with constructs expressing DIP-IRES mCherry or co-expressed with a Dpr-IRES mCherry construct. One-two days post transfection cells were counted and diluted to 1×10^6^ cells/mL in 1x PBS 0.2% BSA, pH 7.4. 45 μL of cells were then added to 96 well plates (Thermo Scientific 262162), and 5 μL of Dpr-Fc protein diluted in 1x PBS 0.2% BSA at 10 times the desired concentration was added. 96 well plates were then incubated on an Elexa E5 Platform Shaker (New Brunswick Scientific) for 45 minutes at room temperature. Cells were then washed three times by centrifuging and removing supernatant, and then incubated with an anti-rat secondary antibody (ThermoFisher, A-21247) diluted 1:300 in 1x PBS 0.2% BSA, and again incubated for 30 minutes shaking, washed 3 times, and then analyzed by flow cytometry on a Novocyte Quanteon (Agilent) or SH800S Cell Sorter (SONY). Gated live cells were sub-gated for mCherry or mCherry/GFO, and GFP-positive cells were sub-gated for Alexa-647. Analysis of flow data was done in FlowJo (FlowJo LLC).

### Computational protocol for GPI-anchor site prediction

Theoretical predictions regarding the mode of membrane attachment in this neuronal protein family have been elusive^26^. Notably, only a handful of family members, including Dpr12, DIP-δ, and DIP-α were predicted to possess a GPI-anchor site (ω-site) when using the default settings of prediction programs such as NetGPI1.1^43^ and PredGPI^44^, including Dpr12, DIP-δ, and DIP-α. Many DIP/Dpr family members, including Dpr6, Dpr8, Dpr10, and DIP-β, were not computationally predicted to have GPI sites.

The ω-site, which designates the location where the protein is cleaved and subsequently connects with the GPI anchor, is traditionally believed to be primarily located near the C-terminus. However, while DIPs and Dprs lack detectable GPI motifs at their C-terminal regions, experimental data suggests that they are indeed GPI-anchored^45^. The ω-site is defined by a distinct amino acid sequence (highlighted in red in Fig. S6A) that is identified by the transamidase complex. Following the ω-site, GPI-anchored proteins typically exhibit a short hydrophilic region (illustrated in blue) succeeded by 10-20 hydrophobic amino acids (depicted in orange). This sequence acts as the GPI attachment motif and is removed during the GPI attachment process.

In order to improve GPI anchor site prediction we developed an approach premised on the idea that the GPI motif might not always be positioned proximal to the C-terminus. Our step-by-step procedure for ω-site prediction, illustrated using DIP-β as an example, is depicted in **Supplementary Figure S5**. First, we generate truncated versions of the protein. This involves systematically removing one residue at a time from the C-terminal tail (see Fig. S5A). All the truncated sequence versions are then batch submitted to the prediction webserver at NetGPI-1.1^43^. Next, we examine the batch submission’s tabulated results to pinpoint the ω-site’s position. Using DIP-β as a case in point: the full-length wild-type (WT) protein, comprising 555 amino acids (aa), is not predicted as GPI-anchored. However, its truncated versions, ranging from 477 to 499 aa in length, are. These truncated forms consistently identify the ω-site at Ser458 (refer to Fig. S5B). In the WT DIP-β sequence, the NetGPI probability for the ω-site reveals a faint signal at S458. Yet, this signal isn’t strong enough for the algorithm to register a positive prediction (see Fig. S5C). In contrast, the truncated versions exhibit a significantly amplified NetGPI probability signal (as highlighted in Fig. S5D).

Our initial protocol did not yield successful identification of GPI-anchor sites in Dpr10. Therefore, we adopted a different algorithm, PredGPI^44,46^, to analyze the truncated versions of the Dpr10 protein (see **Supplementary Data File 2**). Instead of a consensus prediction, the ω-site was identified within a sequence range from W320 to G325. It’s noteworthy that similar patterns of multiple potential sites were observed across various *Drosophila* species.

In various fruit flies, the GPI motifs of selected DIPs and Dprs are annotated in the multiple sequence alignments (MSAs) presented in Fig. S6B. The ω-sites, highlighted in red, were discerned through the computational method described earlier. Meanwhile, the potential hydrophobic region is emphasized with an orange underline (see Fig. S6B). MSAs were performed on the orthologous sequences^11^ of DIPs and Dprs using Clustal Omega^47^ and visualized with Jalview^48^. While multiple isoforms exist for each DIP/Dpr protein, only the longest isoforms were incorporated into the MSAs. *Drosophila* species with only intermediate sequencing coverage^49^ — insufficient to consistently discern all protein isoforms or determine their full length — were omitted from our analysis.

### Drosophila melanogaster rearing and strains

All fly strains were reared under standard laboratory conditions at 25°C on molasses or cornmeal containing food. Males and females were chosen at random. The relevant developmental stage is adult, which refers to 3-5 days post eclosion. UAS-DIP-δ, UAS-Dpr12 and the dpr12 mutant allele Δ50-81 were all previously generated by the Schuldiner lab^14^. R71G10-GAL4 on the 2nd chromosome was also generated by the Schuldiner lab^16^. UAS-Dpr10, DIP-α-Gal4 and UAS-Dpr10 RNAi and Dpr6 RNAi were previously described^6,9,12–15^. All genetic experiments were conducted using Dpr10 isoform D.

### Dissection, immunostaining and microscopy

For adult leg dissection, flies were first immersed in 70% Ethanol for ∼1 min and rinsed in 0.3% Triton-X in PBS 3X. Abdomen and heads were then removed and legs attached to thorax were fixed overnight at 4C in 4% paraformaldehyde in 0.3% Triton-X in PBS. The next day, legs and thorax were washed five times in 0.3% Triton-X in PBS then stored in 80% VECTASHIELD (in PBS) overnight at 4C. Legs were then removed from the thorax and placed onto a glass slide in a drop of VECTASHIELD mounting medium (Vector Labs). 18X18mm coverslips were then placed on top with dental wax on the corners to prevent the coverslip from crushing the legs. 0.5 µm-thick sections in the Z axis were imaged using a Zeiss LSM 800 Confocal Laser Scanning Microscope.

For MB imaging, the brains of adult flies were dissected in cold ringer solution, fixed using 4% paraformaldehyde (PFA) for 20 minutes at room temperature (RT), and then washed in Phosphate Buffer with 0.3% Triton-X (PBT; 3 x immediate washes followed by 3 x 20-minute washes). Non-specific staining was blocked using 5% heat inactivated goat serum in PBT, and brains were then subjected to primary antibody staining overnight at 4°C. Primary antibodies included chicken anti-GFP 1:500 (AVES) and mouse anti-FasII 1:25 (1D4; DSHB). Brains were then washed with PBT (3 x immediate washes followed by 3 x 20-minute washes), stained with secondary antibodies for 2 hours at RT, and washed again. Secondary antibodies included FITC Goat anti-chicken and Alexa fluor 647 goat anti-mouse, both used at 1:300 (Invitrogen). The brains were mounted on Slowfade (Invitrogen) and imaged with Zeiss LSM 980 confocal microscope. Images were processed with ImageJ (NIH).

### Motor Neuron and MB quantification and statistical analysis

To quantify leg motor neurons only one T1 leg from each animal was imaged and their neurons traced using the Simple Neuron Tracer from ImageJ. For each neuron, with the exception of the αTi-tadm, the tracing began at the first bifurcation point. For the αTi-tadm, only the most distal branch was traced due to presence of collateral branches which would have potential misrepresented the terminal branch number and branch length. Both total terminal branch number and the sum of cable length values were separately plotted and analyzed using the GraphPad Prism 9.0 software. For all graphs statistical significance was determined using an unpaired nonparametric two tailed Mann-Whitney test. Error bars represent mean with 95% confidence intervals. ns= no statistical difference. *p<0.05 **p<0.01 ***p<0.001 ****p<0.0001.

To quantify MB phenotypes (**Fig. 6F**), blind ranking of γ4/5 defect severity was performed by an independent investigator, on a scale of 1 (strong defect) to 4 (WT-like morphology), as demonstrated in Bornstein et al. 2021^14^ (Fig. EV1D). The 4 groups were compared by Kruskal-Wallis test, followed by pairwise Mann-Whitney U test corrected for multiple comparisons using FDR.

### Generation of *DIP-α^short^* Allele using CRISPR

We chose two protospacer sequences that were ∼375 bp away from each other, in the exon region coding for the DIP-α membrane linker, to create a large 123 amino acid deletion. The deletion was replaced with a sequence coding for Glycine-Glycine-Serine (GGS), consistent with cloning in cell culture experiments. High score protospacer sequences were chosen on http://crispr.dfci.harvard.edu/SSC/. We cloned both protospacers into a pCFD5_w plasmid (Addgene 112645) and co-injected the plasmid and single-stranded repair template (Integrated DNA Technologies) into CAS0001 flies (Rainbow Transgenic Flies). Injected embryos were crossed with balancer lines, and screened in F1 for individual flies carrying the mutation. A mutant stock was established from this single F1. sgRNA and ssODN sequences are listed below. Detailed protocols are available upon request.

DIP-α^short^ deleted sequence: AACGGCGGCGGAAAAGGAGGTGGAGCGGGCGGAAGCCTGGATGCGGATGCCAATGACATTTTGAAGCAGAAACAACAAGTCAAAGTCACTTATCAGCCGGAGGACGAGGAGCTGCAGTACGGATCCGTCGAGGATTTCGAGGCGGAAGGCGGCGAGGGCGGGGGCCTGACGCCGTTATCGCCGCACGTTTACTACACCAGCGGCAATAAACCGGCCACACATAAGCCGGGCAACTCCGGCGGCAATCAGCATCTACATCAGCAGCACCACCATCATCACCACCACAACAACAACAACAACAACCAGCAACACAGTGGTGGGCCAGGTGGTGGCCCAACTGGCGGTGATGCGGGTTCACTTGGTGGCGAA

Insertion replacing deleted sequence: GGTGGAAGC

sgRNA 1: CGTAATAAAAATCCGCTGAA

sgRNA 2: TCACTTGGTGGCGAAATGGG

ssODN (Repair Template):

CCCGCTGCCCTACAGAAATCCCCGGACCGAATCGTAATAAAAATCCGCTGGGTGGAAGCATGGGAGGCATCACCCGCAAGCCACCGCCATATTATGGTGGCAATACGGAAGTGCGAGGC

### Statistical Information

All data were analyzed with GraphPad Prism 5.0 software (San Diego, CA, USA).

## Author Contributions

N.C.M., D.H.L., H.M., A.P.S., O.S., R.P.M. B.H. and L.S. conceptualized the project, designed experiments, and analyzed data. N.C.M. P.S.K. and F.P. conducted biochemistry experiments. D.H.L. and K.D.R. conducted NMJ experiments. H.M. and B.B. conducted MB experiments. A.P.S. designed and conducted computational experiments. J.E.B. provided reagents. H.G. conducted RNA seq experiments. K.D.R. generated the CRISPR/Cas9 allele. O.S., R.S.M., B.H., and L.S., designed experiments, analyzed data, supervised experiments, and provided funding. N.C.M., R.S.M., B.H. and L.S. wrote the draft. N.C.M., D.H.L., H.M., A.P.S., K.D.R., O.S., R.S.M., B.H., and L.S. edited and revised the manuscript. All authors commented on and approved the final draft.

## Acknowledgements

This work was supported by NSF grant MCB-1914542 (B.H.) and NIH R01NS070644 (R.S.M.), European Research Council (ERC) advanced grant #101054886 ‘‘NeuRemodelBehavior’’ to O.S. O.S. is an incumbent of the Prof. Erwin Netter Professorial Chair of Cell Biology. We thank Lalanti Venkatasubramanian for images in the Supplementary Figures.

## Supplementary

**Figure S1. Related to Figure 1:**
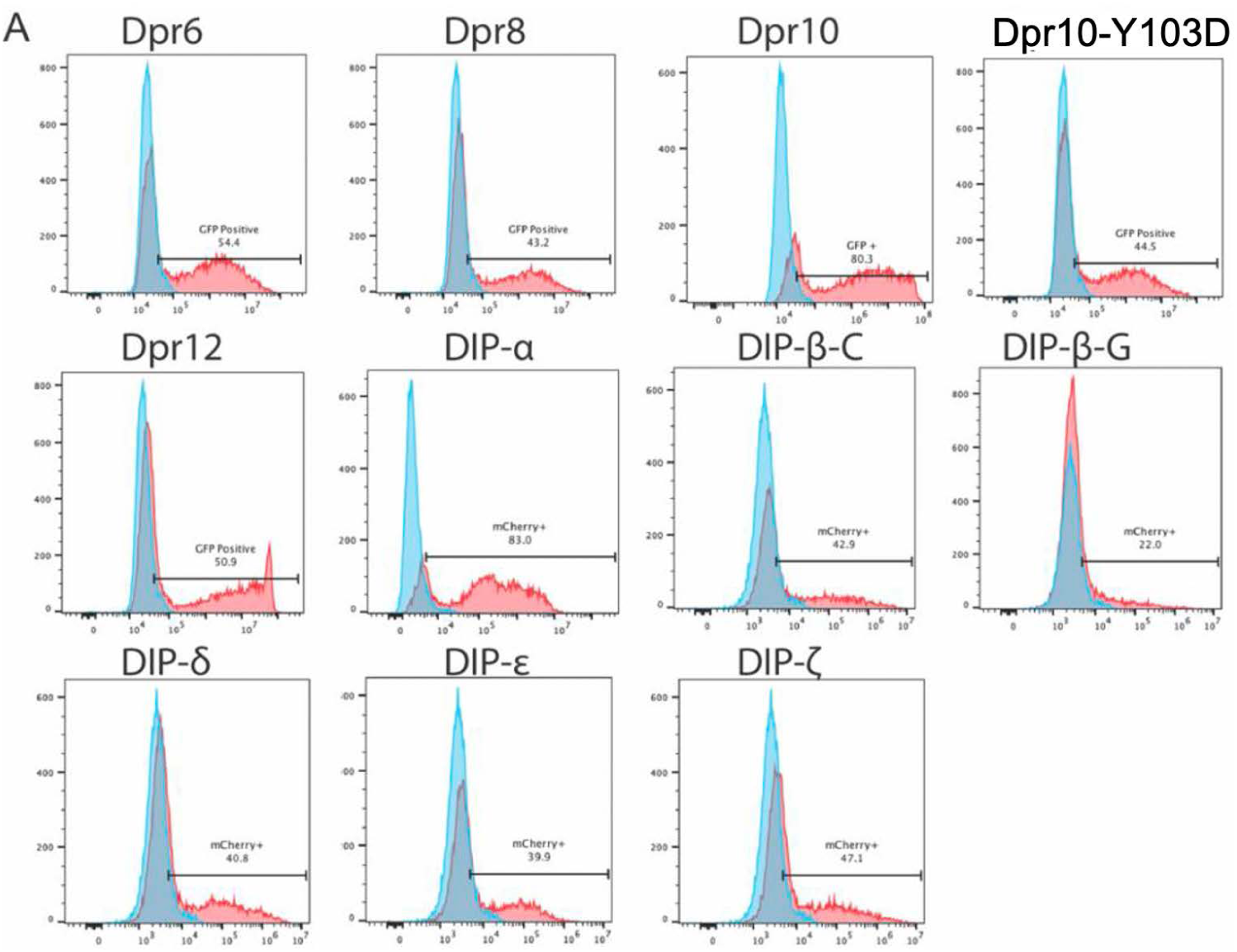
mCherry and GFP expression of full lenth DIP/Dpr construct. **(A**) Percent positive of mCherry (DIPs) and GFP (Dprs) expressed as full length proteins with an IRES tag in mammalian cells.

**Figure S2. Related to Figure 1:**
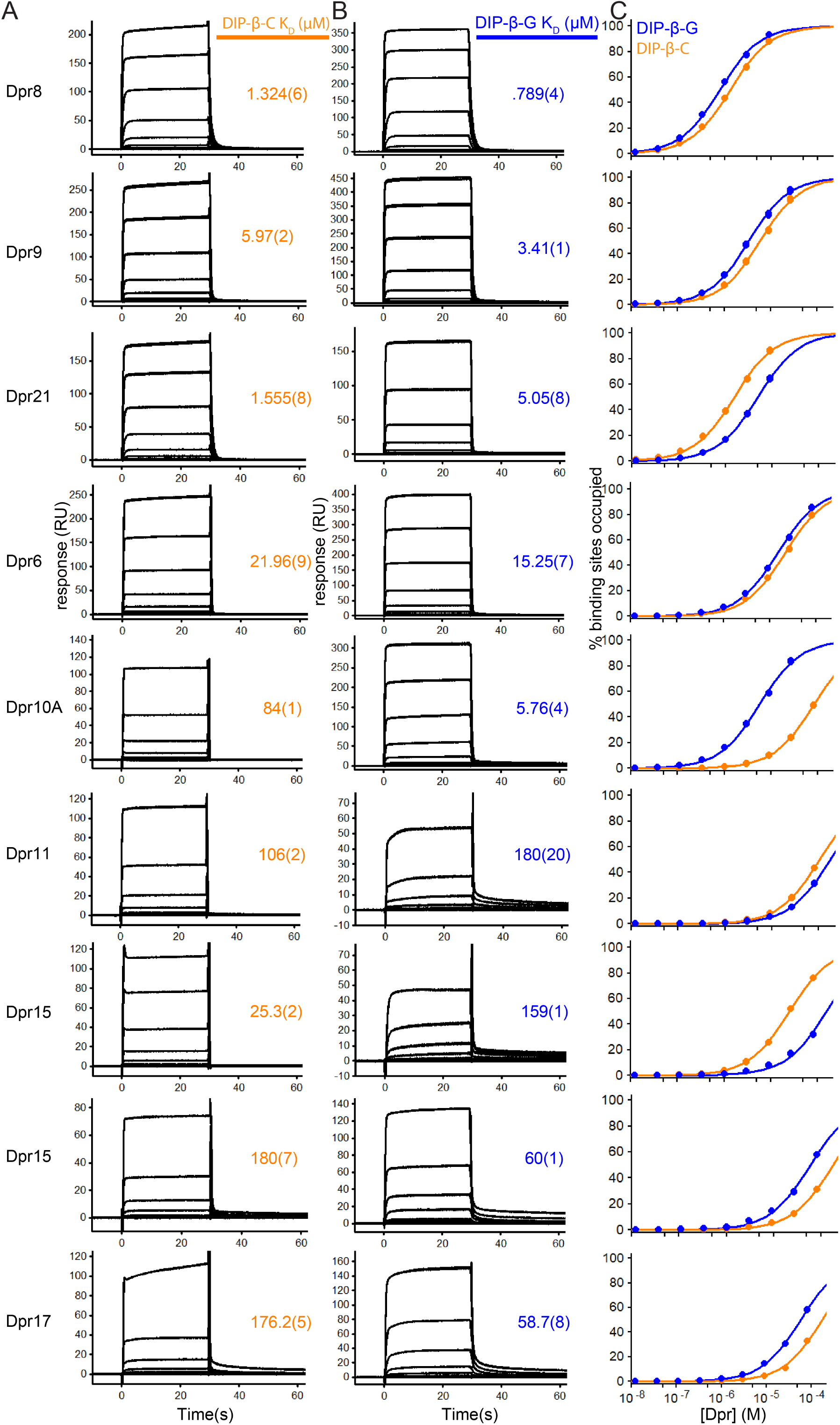
Differential K_D_s of DIP-β C and G isoforms for Dpr10. **(A)** Sensograms of Dpr analytes binding over DIP-β-G immobilized surface and **(B)** Sensograms of Dpr analytes binding over DIP-β-G immobilized surface **(C)** The fit of the binding data to 1:1 binding isotherms to calculate K_D_.

**Figure S3. Related to Figure 2:**
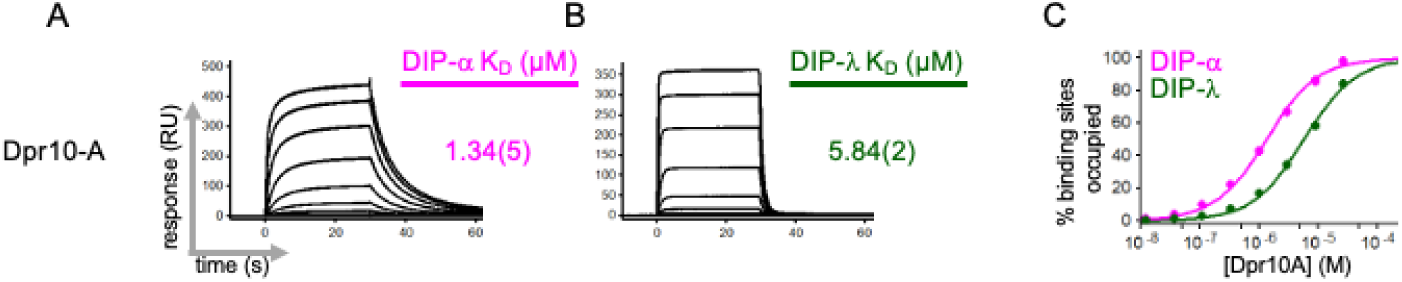
K_D_ of Dpr10-A for DIP-ɑ. **(A)** Sensograms of Dpr10-A analytes binding over DIP-ɑ immobilized surface **(B)** Sensograms of Dpr10-A analytes binding over DIP-λ immobilized surface **(C)** The fit of the binding data to 1:1 binding isotherms to calculate K_D_.

**Figure S4. Related to Figure 2:**
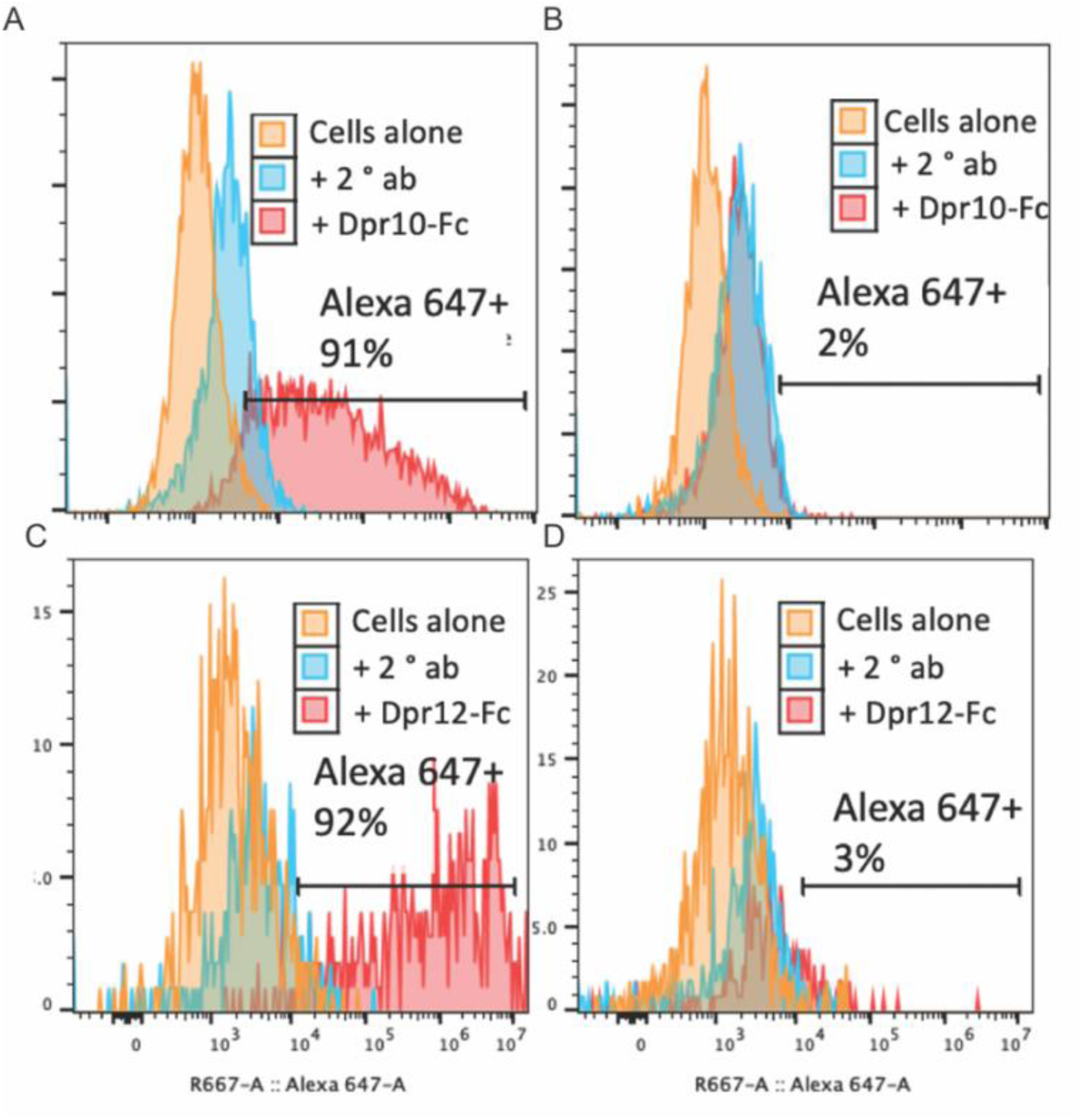
*Cis* inhibition occurs between certain DIP::Dpr pairs Flow Plot. **(A)** Representative flow plots showing binding differences of 300 nM Dpr10-Fc to DIP-ɑ expressing cells or **(B)** DIP-ɑ/Dpr10 co-expressing cells. **(C)** Representative flow plots showing binding differences of 300 nM Dpr12-Fc to DIP-**δ** expressing cells or **(D)** DIP-**δ**/Dpr10 co-expressing cells.

**Figure S5. Related to Figure 4:**
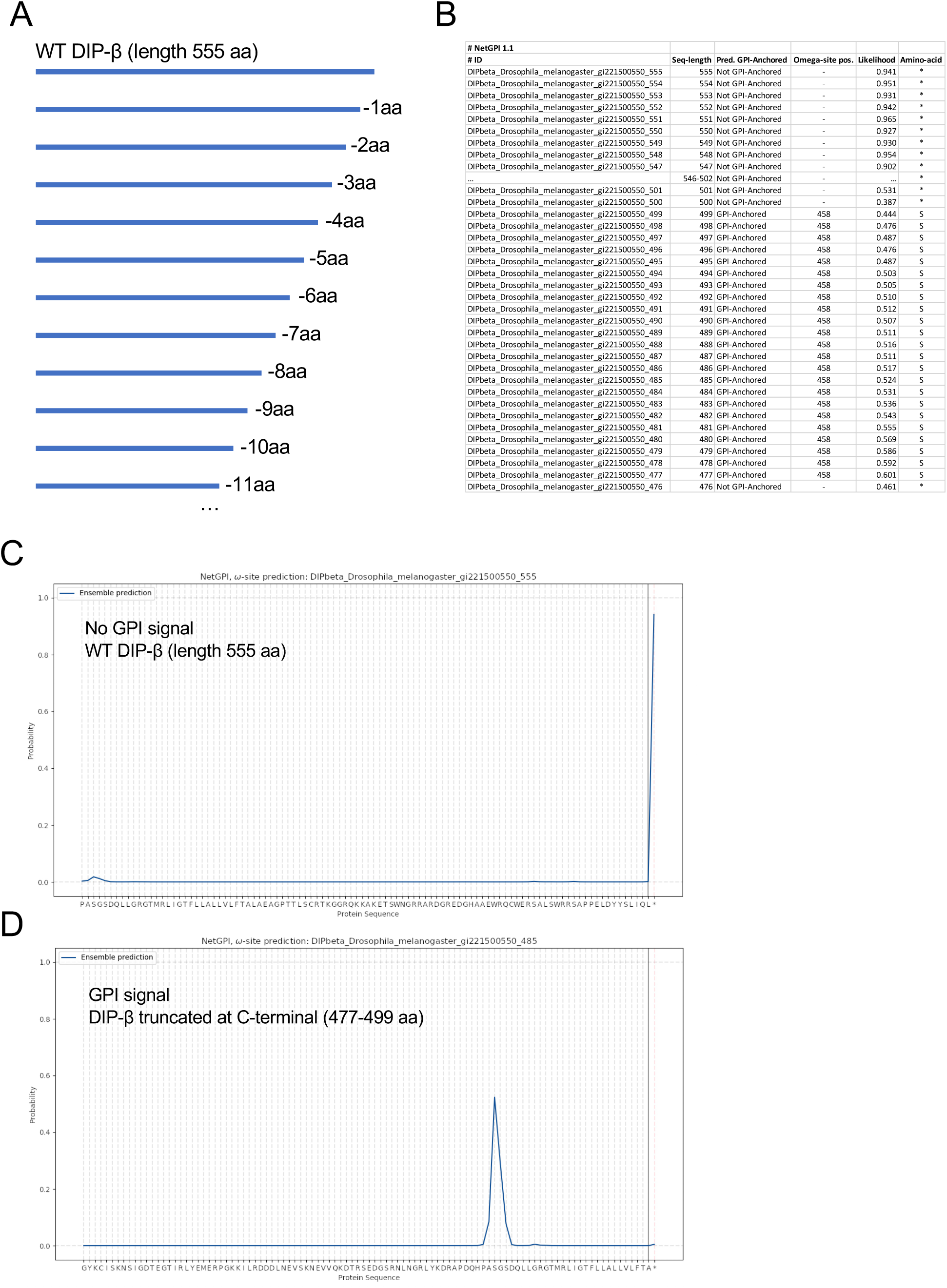
Protocol of GPI ω-site prediction on an example of DIP-β. **(A)** Schematic demonstrating single amino acid deletion method of searching for GPI anchor. **(B)** Representative output for GPI anchor search. **(C-D)** Net GPI outputs for different DIP-β inputs.

**Figure S6: Related to Figure 4:**
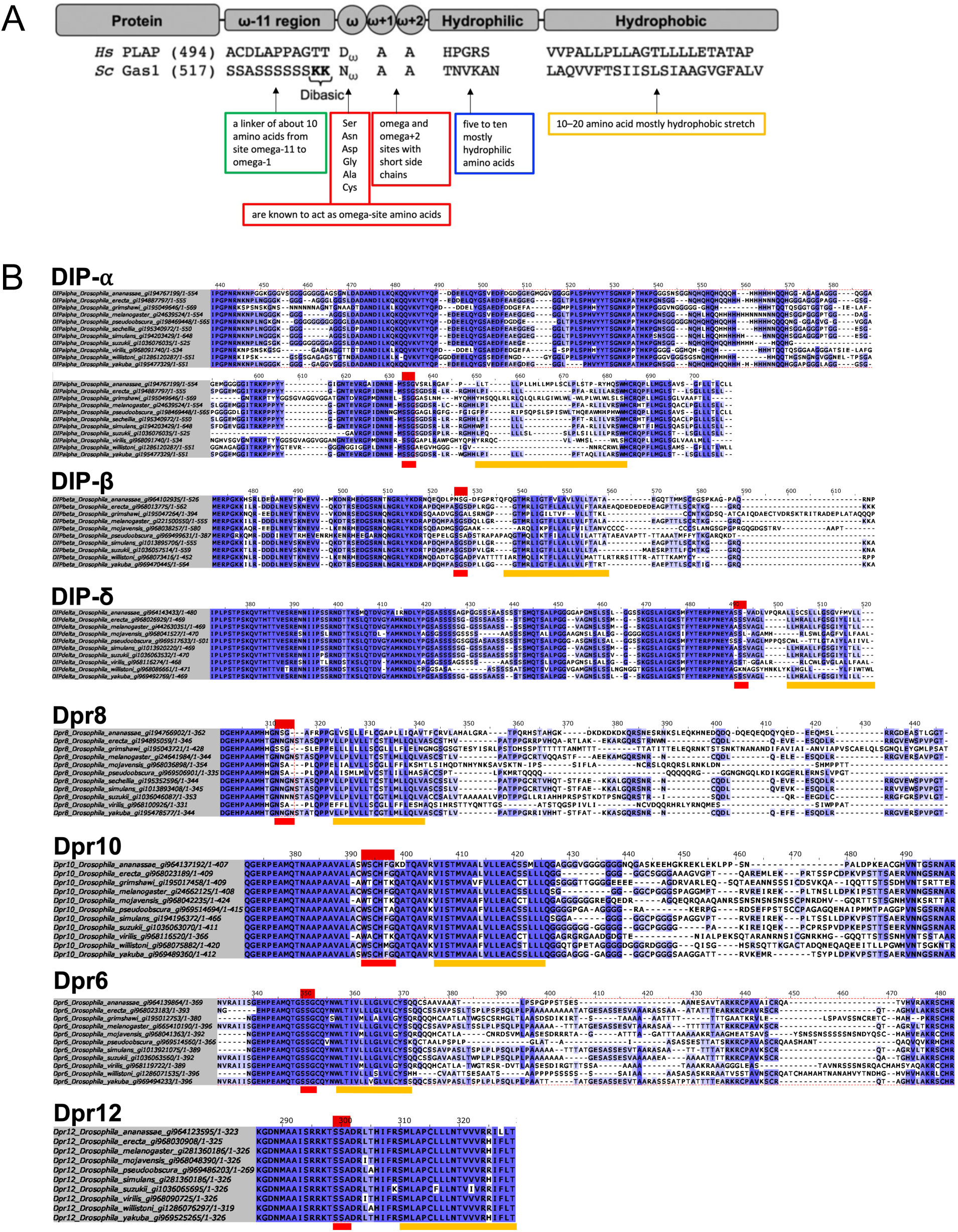
GPI sequence signatures. **(A)** Anticipated amino acid characteristics of the GPI-anchored protein. **(B)** Multiple sequence alignment (MSA) of selected DIP and Dpr C-terminal regions across various Drosophila species; ω-sites highlighted in red and the potential hydrophobic region in orange. Alignment executed via Clustal Omega and visualization achieved with Jalview. The coloring in the MSA reflects sequence identity, with deeper shades of blue indicating higher levels of sequence conservation.

**Figure S7. Related to Figure 6:**
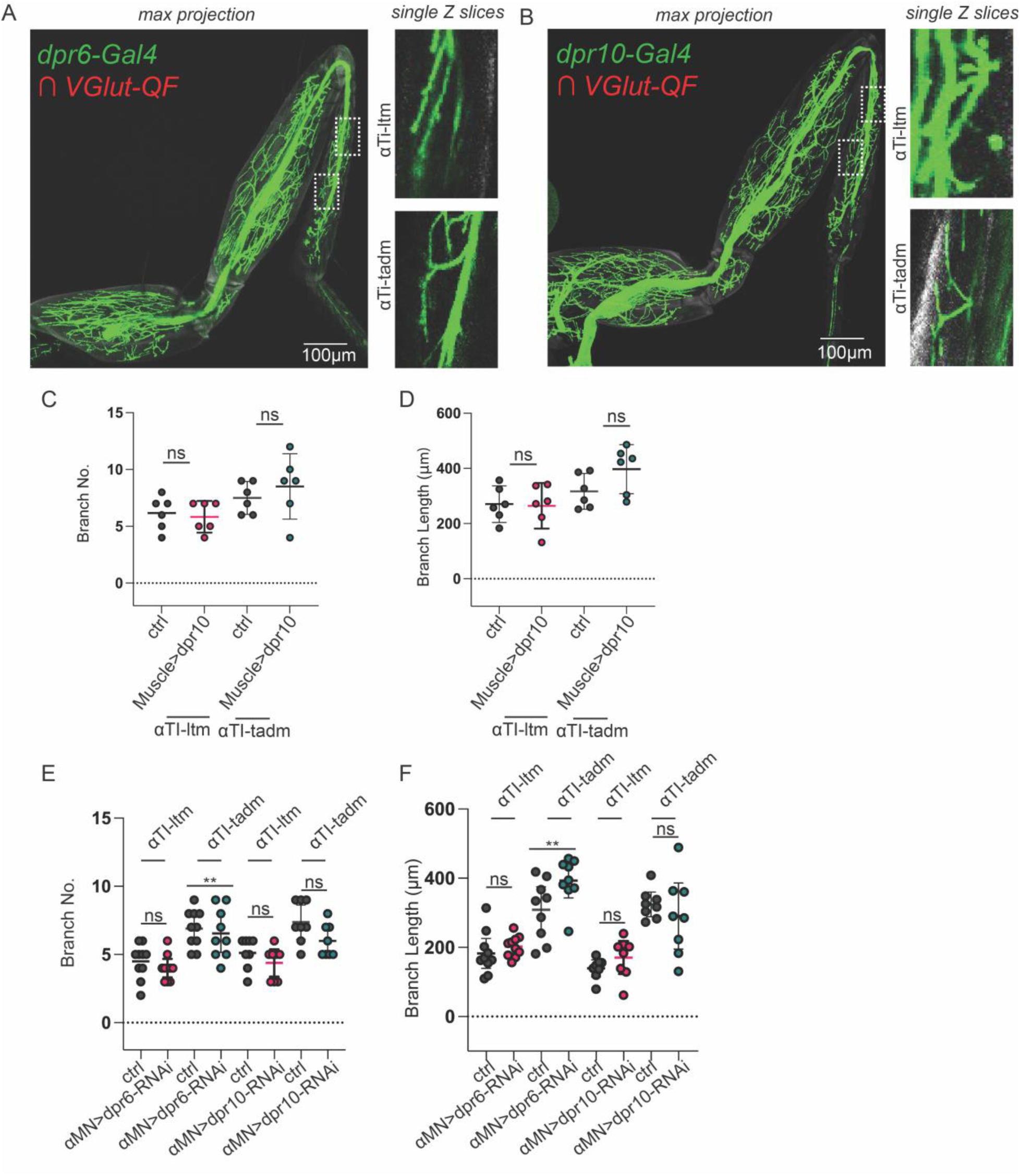
Expression Pattern of dpr6 and dpr10 in MNS replicated from Venkatasubramanian et al., and KD/Overexpression Controls. dpr6-T2A-Gal4 **(A)** and dpr10-T2A-Gal4 **(B)** is restricted to glutamatergic MNs in the T1 legs using a genetic intersectional approach. **(C)** Quantification of the number of αTi-ltm and αTi-tadm terminal branches. Ctrl, *DIP-α-T2A-QF>>10XUAS6XGFP.* αTi-ltm n= 6. αTi-tadm n=6 *Muscle>dpr10, DIP-α-T2A-QF>>10XUAS6XGFP, Mef2-Gal4>UAS-dpr10.* αTi-ltm n=6. αTi-tadm n=6 **(D)** Quantification of the sum of αTi-ltm and αTi-tadm terminal branch length. Ctrl, *DIP-α-T2A-QF>>10XUAS6XGFP.* αTi-ltm n= 6. αTi-tadm n=6 *Muscle>dpr10, DIP-α-T2A-QF>>10XUAS6XGFP, Mef2-Gal4>UAS-dpr10.* αTi-ltm n=6. αTi-tadm n=6. **(E)** Quantification of the number of αTi-ltm and αTi-tadm terminal branches. Ctrl, *DIP-α-T2A-Gal4>>20XUAS6XGFP.* αTi-ltm n= 10. αTi-tadm n=10 *αMN>dpr6-RNAi, DIP-α-T2A-Gal4>>20XUAS6XGFP,UAS-dpr6-RNAi.* αTi-ltm n=10. αTi-tadm n=10 Ctrl, *DIP-α-T2A-Gal4>>20XUAS6XGFP.* αTi-ltm n= 8. αTi-tadm n=8 *αMN>dpr10-RNAi, DIP-α-T2A-Gal4>>20XUAS6XGFP,UAS-dpr10-RNAi.* αTi-ltm n=8. αTi-tadm n=8. **(F)** Quantification of the sum of αTi-ltm and αTi-tadm terminal branch length. Ctrl, *DIP-α-T2A-Gal4>>20XUAS6XGFP.* αTi-ltm n= 10. αTi-tadm n=10 *αMN>dpr6-RNAi, DIP-α-T2A-Gal4>>20XUAS6XGFP,UAS-dpr6-RNAi.* αTi-ltm n=10. αTi-tadm n=10 Ctrl, *DIP-α-T2A-Gal4>>20XUAS6XGFP.* αTi-ltm n= 8. αTi-tadm n=8 *αMN>dpr10-RNAi, DIP-α-T2A-Gal4>>20XUAS6XGFP,UAS-dpr10-RNAi.* αTi-ltm n=8. αTi-tadm n=8.For all graphs statistical significance was determined using an unpaired nonparametric two tailed Mann-Whitney test. Error bars represent mean with 95% confidence intervals. ns= no statistical difference. *p<0.05 **p<0.01 ***p<0.001 ****p<0.0001.

**Figure S8.**
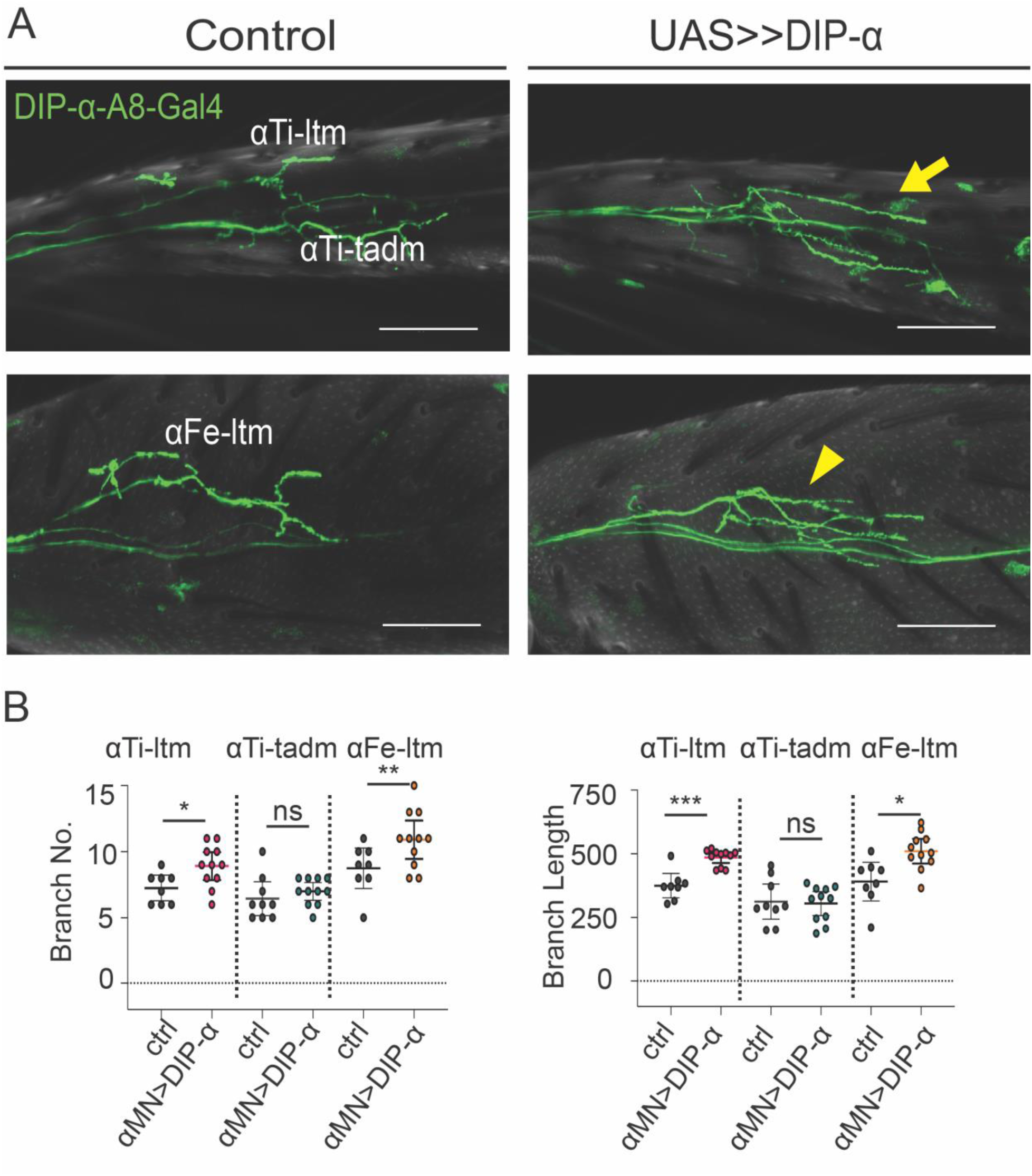
Overexpressing DIP-α in MNs causes αMN axons to increase in length. **A)** Representative Images of *DIP-α-A8-Gal4>>20XUAS6XGFP controls, and DIP-α-A8-Gal4>>20XUAS6XGFP, UAS-DIP-α.* Yellow arrow indicates longer αTi-ltm axons. Yellow arrowhead indicates longer αFe-ltm axons. Scale bar= 50µm **B)** Quantification of the number of αTi-ltm, αTi-tadm and αFe-ltm terminal branches.Ctrl, *DIP-α-A8-Gal4 20XUAS6XGFP.* αTi-ltm n= 8. αTi-tadm n=9 αFe-ltm n=8. *DIP-α-A8-Gal4>>20XUAS6XGFP, UAS-DIP-α.* αTi-ltm n=11. αTi-tadm n=11 αFe-ltm n=11. Quantification of the sum of αTi-ltm, αTi-tadm, and αFe-ltm, branch length. Ctrl, *DIP-α-A8-Gal4 20XUAS6XGFP.* αTi-ltm n= 8. αTi-tadm n=9 αFe-ltm n=8. *DIP-α-A8-Gal4>>20XUAS6XGFP, UAS-DIP-α.* αTi-ltm n=11. αTi-tadm n=11 αFe-ltm n=11.For all graphs statistical significance was determined using an unpaired nonparametric two tailed Mann-Whitney test. Error bars represent mean with 95% confidence intervals. ns= no statistical difference. *p<0.05 **p<0.01 ***p<0.001*. Ctrl-Control, MN-motor neuron*.

