## Supplementary Data File 1 for "*Cis* inhibition of co-expressed DIPs and Dprs shapes neural development"

Supplemental Data File I

|  | DIP-α | DIP-β-C | DIP-β-G | DIP-δ | DIP-ε | DIP-ζ |
| --- | --- | --- | --- | --- | --- | --- |
| Dpr6 | 2.06 | 19.4 | 15.25 | >300 | 210 | 151 |
| Dpr8 | >500 | 1.52 | .789 | >500 | >1000 | >500 |
| Dpr10-A | 1.34 | 84 | 5.76 | unknown | unknown | unknown |
| Dpr10-D | 1.67 | 54.9 | 35.9 | 218 | >1000 | >500 |
| Dpr12 | >1000 | >500 | unknown | 2.44 | >1000 | >500 |

Table I- K_D_s important for this study

K_D_s are indicated in micromolar and appropriate references are shown below

Reported in this study (Supplemental Figures S2 and S3)

Reported in Cosmanescu, et. al, 2018

Reported in Carrillo, et. al, 2015
