## Supplementary figures and images for "*Cis* inhibition of co-expressed DIPs and Dprs shapes neural development"

### Supplementary Data File 2

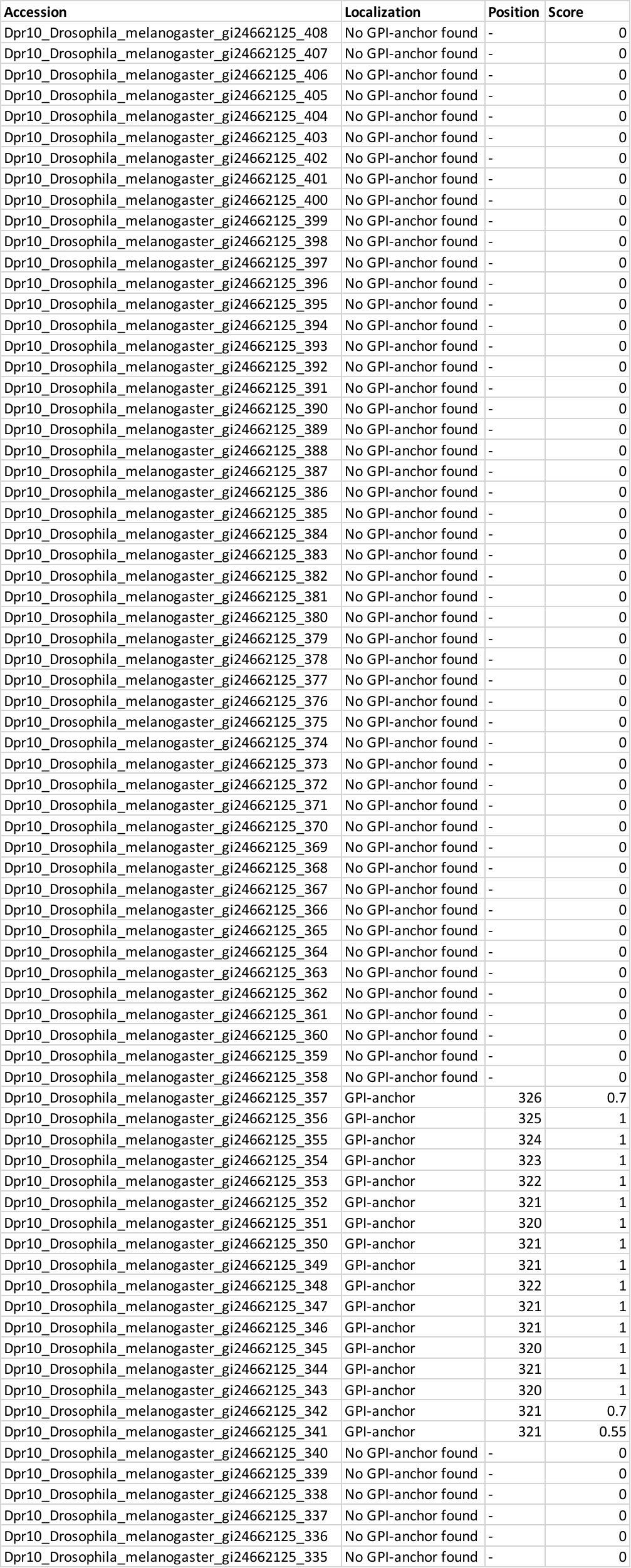
